# A dual mechanism of enhancer activation by FOXA pioneer factors induces endodermal organ fates

**DOI:** 10.1101/2020.08.28.263020

**Authors:** Ryan J. Geusz, Allen Wang, Dieter K. Lam, Nicholas K. Vinckier, Konstantinos-Dionysios Alysandratos, David A. Roberts, Jinzhao Wang, Samy Kefalopoulou, Yunjiang Qiu, Joshua Chiou, Kyle J. Gaulton, Bing Ren, Darrel N. Kotton, Maike Sander

**Author notes:** These authors contributed equally (Ryan J. Geusz, Allen Wang).

## Abstract

FOXA pioneer transcription factors (TFs) displace nucleosomes and prime chromatin across enhancers of different endodermal organs in multipotent precursors before lineage induction. Here, we examined patterns and mechanisms of FOXA target site engagement using human pluripotent stem cell models of endodermal organ development. Unexpectedly, we find that only a subset of pancreatic, hepatic, and alveolar enhancers are FOXA-primed, whereas the majority are unprimed and engage FOXA only upon lineage induction. Analysis of sequence architecture revealed more abundant and stronger FOXA motifs at primed than unprimed enhancers and enrichment for lineage-specific TF motifs at unprimed enhancers. We show that FOXA recruitment to unprimed enhancers specifically depends on lineage-specific TFs, suggesting that regulatory DNA sequence logic governs temporal FOXA recruitment. Our findings suggest that FOXA-mediated enhancer priming broadly facilitates initiation of organ lineage programs, while secondary FOXA recruitment by lineage-specific TFs to the majority of enhancers confers organ specificity to gene expression.

## INTRODUCTION

The pancreas, liver, and lung develop from the foregut endoderm in response to local signaling cues that specify lineage identity by inducing organ-specific gene expression. The competence of organ lineage precursors to activate lineage-specific genes in response to inductive signals is acquired during endoderm development (Wang et al., 2015; Zorn and Wells, 2009). Coincident with the acquisition of competence, the transcription factors (TFs) FOXA1 and FOXA2 (henceforth abbreviated FOXA1/2) are recruited to enhancers of foregut-derived organ lineages, leading to a gain in chromatin accessibility and H3K4me1 deposition (Genga et al., 2019; Lee et al., 2019; Wang et al., 2015), a phenomenon referred to as enhancer priming. Thus, current evidence supports a model wherein FOXA1/2 render foregut endoderm competent to activate organ-specific genes by broadly priming pancreas-, liver-, and lung-specific enhancers before organ-inductive signals trigger enhancer activation and target gene expression. Consistent with this model, studies in model organisms and human pluripotent stem cell (hPSC)-based differentiation systems have shown a requirement for FOXA1/2 in pancreas, liver, and lung development, with the two FOXA TFs functioning in a partially or fully redundant manner (Genga et al., 2019; Lee et al., 2005; Lee et al., 2019; Wan et al., 2005). However, the extent to which the full repertoire of organ lineage-specific enhancers undergoes chromatin priming is unknown.

The mechanisms by which FOXA TFs engage with and open chromatin have been the subject of debate. *In vitro* experiments have shown that FOXA TFs possess pioneering activity, which refers to the specific ability of a TF to engage target sites on nucleosomal DNA and to remodel such regions to increase chromatin accessibility (Cirillo et al., 2002; Iwafuchi-Doi et al., 2016; Soufi et al., 2015). Through their chromatin remodeling activity, FOXA TFs facilitate subsequent binding of other TFs and co-factors that further modify chromatin state and initiate gene expression (Carroll et al., 2005; Cirillo et al., 2002; Gualdi et al., 1996; Hurtado et al., 2011; Magnani and Lupien, 2014; Wang et al., 2009). However, despite their ability to access target sites in closed chromatin *in vitro*, binding site selection of FOXA and other pioneer TFs in cellular contexts has been shown to depend on additional features, such as the local chromatin landscape (Lupien et al., 2008), presence of cooperative binding partners (Caizzi et al., 2014; Donaghey et al., 2018), and strength of the binding motif (Donaghey et al., 2018; Swinstead et al., 2016; Zhang et al., 2015). For example, steroid receptor activation in breast cancer cell lines induces FOXA1 recruitment to sites with degenerate FOXA1 binding motifs (Paakinaho et al., 2019; Swinstead et al., 2016), exemplifying heterogeneity in FOXA target site engagement. The determinants that underlie FOXA binding site selection and FOXA-mediated enhancer priming during cellular transitions of development remain to be explored.

Here, we sought to determine mechanisms of lineage-specific enhancer activation by FOXA1/2 during endodermal organ development, and mapped FOXA1/2 genomic binding throughout a time course of hPSC differentiation into pancreas, liver, or lung alveolospheres. Surprisingly, only a minority of organ lineage-specific enhancers are FOXA1/2-bound prior to lineage induction and exhibit priming, whereas the majority engage FOXA1/2 concomitant with lineage induction. Compared to unprimed organ-specific enhancers, primed enhancers contain DNA sequences more closely matching FOXA consensus motifs and harbor additional sequence motifs for signal-dependent TFs. By contrast, unprimed enhancers are enriched for motifs of lineage-specific TFs and unlike primed enhancers require lineage-specific TFs for FOXA1/2 recruitment. Our findings show that FOXA1/2 regulate foregut organ enhancers through two distinct mechanisms: priming of a small subset of enhancers across organs before lineage induction and activation of a larger cohort of enhancers through cooperative binding with organ lineage-specific TFs. We speculate that the here-identified mechanism of cooperative FOXA1/2 binding with lineage-specific TFs could provide a safeguard against broad activation of alternative lineage programs across different organs.

## RESULTS

### FOXA1 and FOXA2 are necessary for pancreatic lineage induction

To investigate the role of FOXA1/2 in pancreas development, we employed a hPSC differentiation protocol in which cells transition stepwise to the pancreatic fate through sequential exposure to developmental signaling cues (**Figure 1A**). The pancreatic lineage is induced by retinoic acid from gut tube (GT) intermediates, resulting in expression of the pancreatic markers PDX1 in early pancreatic progenitors (PP1) and NKX6.1 in late pancreatic progenitors (PP2). FOXA1 and FOXA2 were expressed from the definitive endoderm (DE) stage onwards (**Figures S1A** and **S1B**), and levels of *FOXA1* and *FOXA2* were similar in GT, PP1, and PP2 (**Figures S1A**).

**Figure 1.**
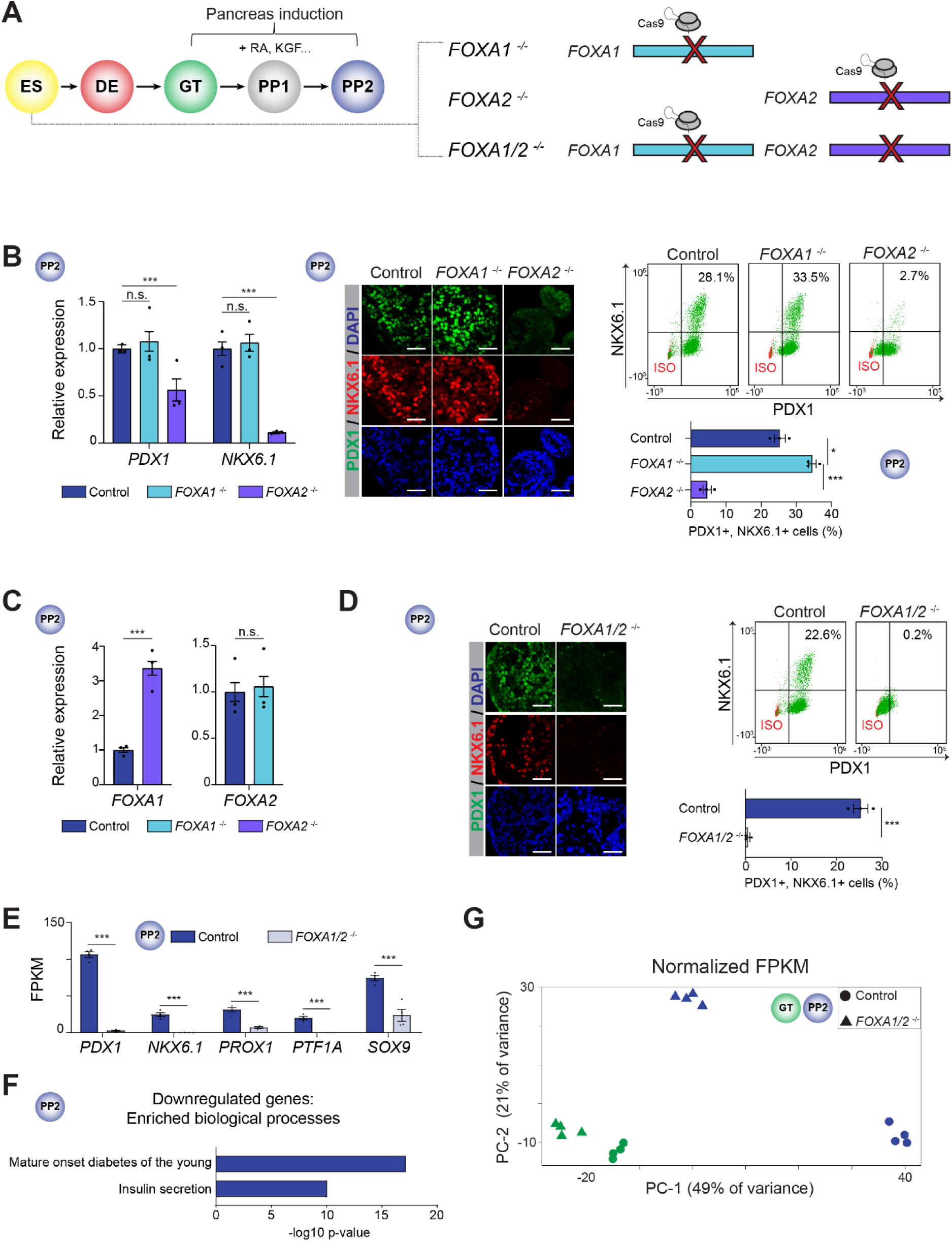
Partially redundant requirement for FOXA1 and FOXA2 in pancreatic lineage induction. (**A**) Schematic of stepwise pancreatic differentiation protocol from hESCs (ES): definitive endoderm (DE), primitive gut tube (GT), early pancreatic progenitor cells (PP1), and late pancreatic progenitor cells (PP2), with indicated genetic modifications in ES. Select growth factors for pancreatic lineage induction are indicated. RA, retinoic acid; KGF, keratinocyte growth factor. (**B**) qPCR analysis of *PDX1* and *NKX6.1* (left), representative immunofluorescent staining (middle), and flow cytometry analysis and quantification of PDX1^+^ and NKX6.1^+^ cells (right) in control, *FOXA1*^-/-^ and *FOXA2*^-/-^ PP2 cells (n = 3 independent differentiations; qPCR: *P* = 0.493, 0.590, 3.12 x 10^-3^, and < 1.00 x 10^-6^ for *PDX1* and *NKX6.1* in control compared to *FOXA1*^-/-^ and *FOXA2*^-/-^ PP2 cells, respectively; flow cytometry: *P* = 1.15 x 10^-2^ and 7.00 x 10^- 4^ in control compared to *FOXA1*^-/-^ and *FOXA2*^-/-^ PP2 cells, respectively; student’s t-test, 2-sided; n.s., not significant). (**C**) qPCR analysis of *FOXA1* and *FOXA2* in control, *FOXA1*^-/-^ and *FOXA2*^-/-^ PP2 cells (n = 3 independent differentiations; *P* = < 1.00 x 10^-6^ and 0.700 for *FOXA1* and *FOXA2* in control compared to *FOXA1*^-/-^ and *FOXA2*^-/-^ PP2 cells, respectively; student’s t-test, 2-sided). (**D**) Representative immunofluorescent staining (left) and flow cytometry analysis and quantification (right) of PDX1^+^ and NKX6.1^+^ cells in control and *FOXA1/2*^-/-^ PP2 cells. (n = 3 independent differentiations; *P* = 2.6 x 10^-3^ in control compared to *FOXA1/2*^-/-^ PP2 cells; student’s t-test, 2-sided). (**E**) mRNA expression levels of pancreatic transcription factors determined by RNA-seq in control and *FOXA1/2*^-/-^ PP2 cells (n = 4 independent differentiations; *P* adj. = 1.08 x 10^-42^, 2.56 x 10^-12^, 4.93 x 10^-20^, 1.00 x 10^-49^, and 2.82 x 10^-4^ for *PDX1*, *NKX6.1*, *PROX1*, *PTF1A*, and *SOX9*, respectively; DESeq2; FPKM, fragments per kilobase per million fragments mapped). (**F**) Enriched gene ontology terms of 2833 downregulated genes (≥ 2-fold decrease, *P* adj. < 0.05) in *FOXA1/2*^-/-^ compared to control PP2 cells. (**G**) Principle component analysis showing variance in total normalized transcriptome between control and *FOXA1/2*^-/-^ cells in GT and PP2. Each plotted point represents one biological replicate. For all qPCR, each plotted point represents the average of three technical replicates. For all immunofluorescence, representative images are shown from n ≥ 2 independent differentiations. Scale bars, 50 µm. For all flow cytometry analyses, representative plots are shown from n = 3 independent differentiations, with isotype control (ISO) for each antibody shown in red and target protein staining in green. See also Figure S1 and Table S1.

To determine a possible requirement for FOXA1 and FOXA2 in pancreas development, we deleted *FOXA1* or *FOXA2* in human embryonic stem cells (hESCs) (**Figures 1A, S1C** and **S1D**) and differentiated control, *FOXA1*^-/-^, and *FOXA2*^-/-^ hESC lines into pancreatic progenitors. Analysis of PDX1 and NKX6.1 expression revealed a requirement for FOXA2 but not FOXA1 for pancreatic lineage induction (**Figure 1B**), consistent with recent findings (Lee et al., 2019). The presence of residual PDX1^+^ and NKX6.1^+^ cells and increased *FOXA1* levels in *FOXA2*^-/-^ pancreatic progenitors (**Figures 1B** and **1C**) suggests FOXA1 partially compensates for FOXA2 deficiency. Therefore, we generated *FOXA1*^-/-^;*FOXA2*^-/-^ (*FOXA1/2*^-/-^) hESC lines (**Figure S1E**) and analyzed phenotypes at the DE, GT, and PP2 stages. At the DE and GT stages, similar numbers of *FOXA1/2*^-/-^ and control cells expressed the DE marker SOX17 and GT marker HNF1B, respectively (**Figures S1F** and **S1G)**, suggesting that absence of FOXA1/2 does not abrogate DE and GT formation. In contrast, pancreas induction was blocked in *FOXA1/2*^-/-^ cells, as evidenced by an almost complete absence of PDX1^+^ and NKX6.1^+^ cells, reduced expression of early pancreatic TFs, and down-regulation (≥ 2 fold change, FDR ≤ 0.05) of genes associated with pancreas-specific biological processes (**Figures 1D-1F** and **Table S1**). Principle component analysis (PCA) of transcriptome data further confirmed that *FOXA1/2*^-/-^ and control cells were similar at the GT stage but differed substantially at the PP2 stage (**Figure 1G**). Together, these findings show that FOXA1 and FOXA2 control pancreatic lineage induction from gut tube lineage intermediates in a partially redundant manner.

### FOXA transcription factors exhibit two temporal patterns of recruitment to pancreatic enhancers

To identify transcriptional targets of FOXA1/2 during pancreatic lineage induction, we mapped FOXA1/2 binding sites at the GT and PP2 stages. Consistent with the partial functional redundancy between FOXA1 and FOXA2 (**Figures 1B-1D**), FOXA1 and FOXA2 binding sites were highly correlated at both stages (**Figure S2A**). FOXA1/2 mostly bound to distal sites (>2.5 kb from TSS; **Figure S2B**), suggesting regulation of enhancers by FOXA1/2. To test this, we defined GT and PP2 enhancers as distal H3K27ac peaks (> 2.5 kb from TSS) and compared enhancer activity based on H3K27ac signal in control and *FOXA1/2*^-/-^ cells at the GT and the PP2 stages. Similar to gene expression (**Figure 1G**), H3K27ac profiles of control and *FOXA1/2*^-/-^ cells were similar at the GT stage but differed significantly at the PP2 stage (**Figure S2C**), showing that *FOXA1/2* deletion has broad impact on regulation of enhancer activity during the GT to PP2 transition.

To investigate specific mechanisms of enhancer regulation by FOXA1/2 during pancreatic lineage induction, we identified all FOXA1/2-bound pancreatic enhancers that are activated upon pancreatic lineage induction. To this end, we first identified enhancers that exhibited a ≥ 2-fold increase in H3K27ac signal from the GT to the PP2 stage (2574 enhancers, hereafter referred to as pancreatic enhancers; **Figures S2D** and **S2E**). As expected, genes near these enhancers were predicted to regulate biological processes associated with pancreas development. Second, we analyzed FOXA1/2 binding at these pancreatic enhancers, revealing that 72% were FOXA1/2-bound at the PP2 stage (**Figure S2F**). Consistent with prior reports (Lee et al., 2019; Wang et al., 2015), we observed FOXA1/2 occupancy at the GT stage preceding pancreatic lineage induction. Surprisingly, however, the percentage of pancreatic enhancers bound by FOXA1/2 was significantly lower at the GT compared to the PP2 stage, implying that not all pancreatic enhancers engage FOXA1/2 before lineage induction. To comprehensively characterize temporal patterns of FOXA1/2 recruitment, we identified all pancreatic enhancers with FOXA1 or FOXA2 binding at the GT and/or PP2 stages and quantified FOXA1/2 ChIP-seq signal at these sites (**Figure 2A**). We observed three distinct patterns of FOXA1/2 occupancy: class I enhancers (561) were bound by FOXA1/2 at both the GT and PP2 stages, class II enhancers (1422) were FOXA1/2-bound only at the PP2 stage, and the overall small group of class III enhancers (118) was FOXA1/2-bound only at the GT stage (**Figure 2A** and **Table S2**). Since the predominant patterns were either maintenance of FOXA1/2 binding (class I) or *de novo* FOXA1/2 occupancy (class II) after pancreas induction, we excluded class III enhancers from further analyses. Analysis of H3K27ac signal intensity at the GT and PP2 stages showed similar patterns of H3K27ac signal at class I and class II enhancers (**Figure 2B**), suggesting that enhancers of both classes are mostly inactive at the GT stage and become activated during pancreatic lineage induction. We identified examples of both class I and class II enhancers in proximity to gene bodies of pancreatic lineage-determining TFs which are expressed in PP2 but not GT, such as *PDX1, HNF1B*, *NKX6.1*, and *MNX1* (**Figure 2C**), and found that activation of both classes of enhancers during the GT to PP2 transition was dependent on FOXA1/2 (**Figure 2D**). Consistent with the H3K27ac pattern, the *PDX1* class I enhancer and the *NKX6.1* class II enhancer are both inactive in GT and active in PP2 in enhancer reporter assays (Wang et al., 2015). Together, this analysis shows that FOXA1/2 recruitment to pancreatic enhancers precedes lineage induction at only a small subset of enhancers, while FOXA1/2 recruitment to the majority of pancreatic enhancers coincides with lineage induction (**Figure 2E**).

**Figure 2.**
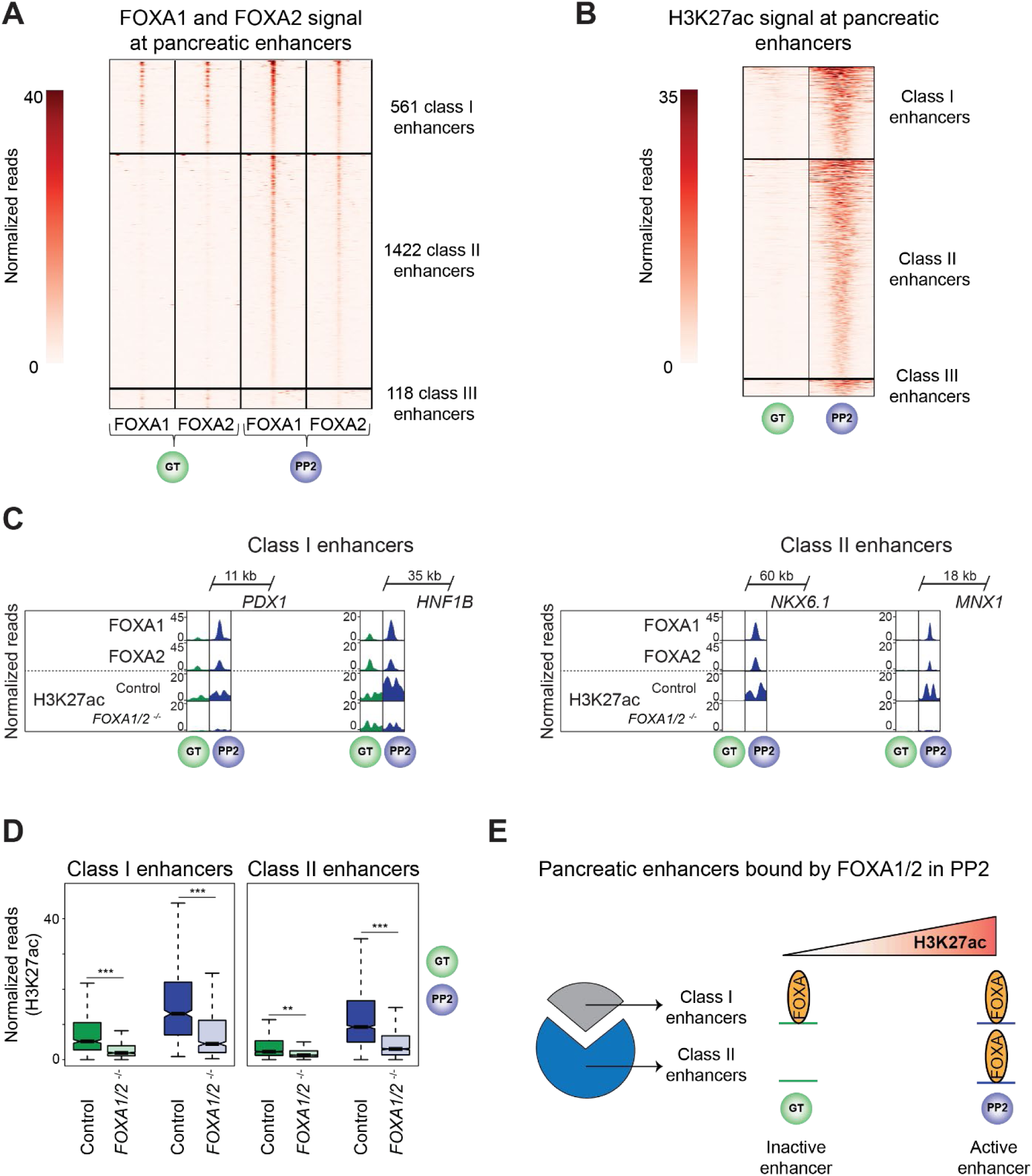
Two distinct temporal patterns of FOXA1 and FOXA2 binding to pancreatic enhancers. (**A** and **B**) Heatmaps showing density of FOXA1 and FOXA2 ChIP-seq reads (**A**) and H3K27ac ChIP-seq reads (**B**) at pancreatic enhancers in GT and PP2. Heatmaps are centered on FOXA1, FOXA2, and H3K27ac peaks, respectively, and span 5 kb. Pancreatic enhancers are classified based on temporal pattern of FOXA1 and FOXA2 occupancy. (**C**) Genome browser snapshots showing FOXA1, FOXA2, and H3K27ac ChIP-seq signal at class I pancreatic enhancers near *PDX1* and *HNF1B* and class II pancreatic enhancers near *NKX6.1* and *MNX1* in GT and PP2. Approximate distance between enhancer and gene body is indicated. (**D**) Box plots of H3K27ac ChIP-seq counts at class I and class II pancreatic enhancers in control and *FOXA1/2*^-/-^ GT and PP2 cells (*P* = < 2.2 x 10^-16^, < 2.2 x 10^-16^, 0.009, and < 2.2 x 10^-16^ for control versus *FOXA1/2*^-/-^ at class I enhancers in GT, class I enhancers in PP2, class II enhancers in GT, and class II enhancers in PP2, respectively; Wilcoxon rank sum test, 2-sided). (**E**) Schematic illustrating the identified pattern of FOXA1/2 occupancy at pancreatic enhancers. All ChIP-seq experiments, n = 2 replicates from independent differentiations. See also Figure S2 and Table S2.

### Primed and unprimed pancreatic enhancers reside in distinct regulatory domains

Given early recruitment of FOXA1/2 to class I but not class II enhancers, we hypothesized that the two classes could differ in their temporal pattern of gain in chromatin accessibility and H3K4me1 deposition, predicting that early FOXA1/2 occupancy at class I enhancers could lead to chromatin priming. As predicted, class I enhancers exhibited open chromatin and H3K4me1 deposition at the GT stage (**Figures 3A, S3A,** and **S3B**). By contrast, class II enhancers acquired these features largely with pancreatic lineage induction (**Figures 3A, S3A,** and **S3B**), identifying primed chromatin as a distinguishing feature of class I enhancers. At both class I and class II enhancers, H3K4me1 deposition and gain in chromatin accessibility during lineage induction was dependent on FOXA1/2 (**Figure 3B** and **S3B**), demonstrating that FOXA1/2 remodel chromatin at both classes of enhancers.

**Figure 3.**
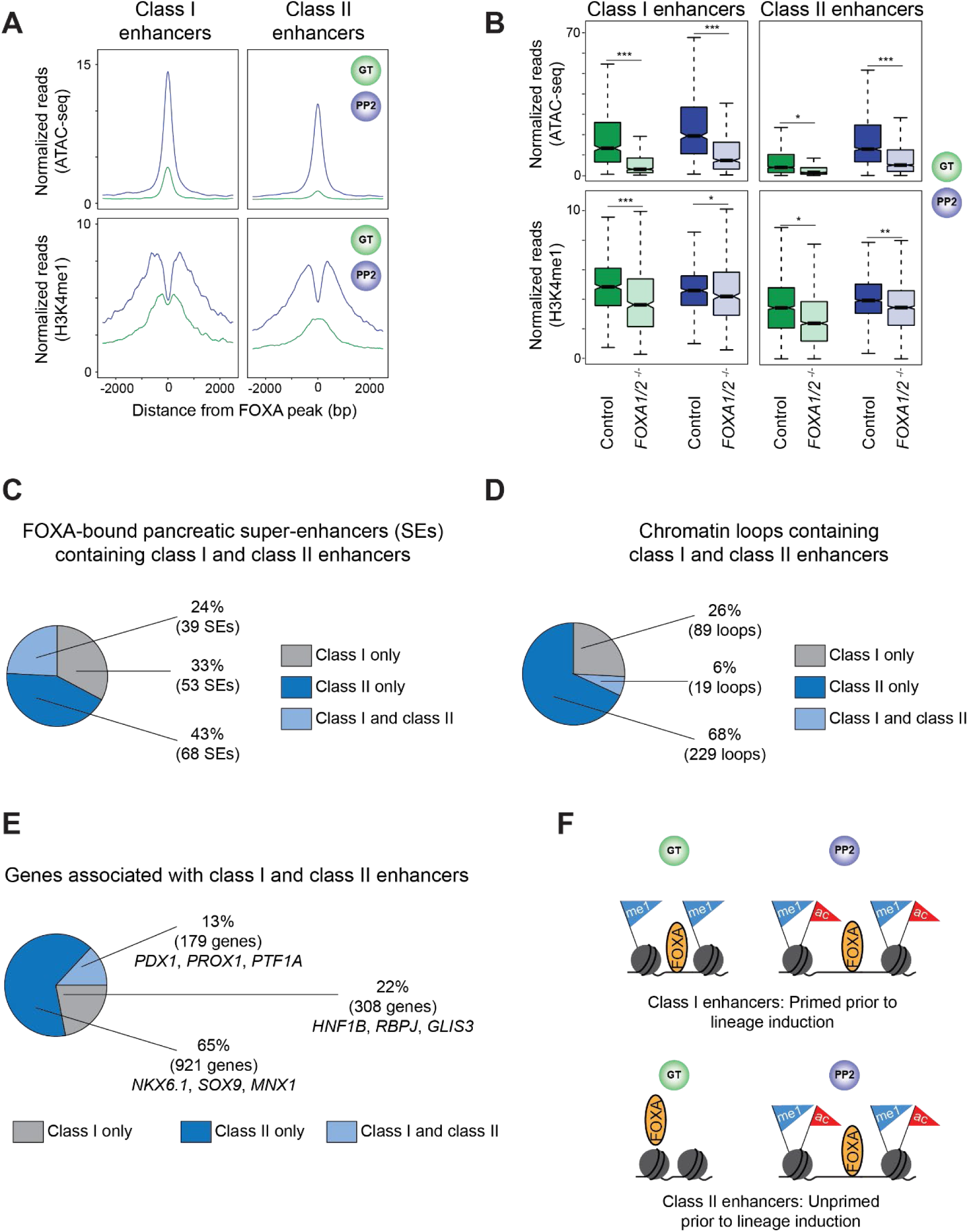
Class I and class II pancreatic enhancers largely map to distinct gene regulatory elements. (**A**) Tag density plots for class I and class II pancreatic enhancers displaying ATAC-seq (top) and H3K4me1 ChIP-seq (bottom) read density in GT and PP2. Plots are centered on FOXA1/2 peaks and span 5 kb. (**B**) Box plots of ATAC-seq (top) and H3K4me1 ChIP-seq (bottom) counts at class I and class II pancreatic enhancers in GT and PP2 for control and *FOXA1/2*^-/-^ cells (*P* = < 2.2 x 10^-16^, < 2.2 x 10^-16^, 0.01, and < 2.2 x 10^-16^ for control versus *FOXA1/2*^-/-^ of ATAC-seq signal at class I in GT, class I in PP2, class II in GT, and class II in PP2, respectively. *P* = < 2.2 x 10^-16^, 0.01, 0.02, and 0.01 for control versus *FOXA1/2*^-/-^ of H3K4me1 signal at class I in GT, class I in PP2, class II in GT, and class II in PP2, respectively; Wilcoxon rank sum test, 2-sided). (**C**) Percentage of FOXA1- and/or FOXA2-bound pancreatic super-enhancers (SEs) in PP2 containing only class I, only class II, or both class I and class II enhancers. (**D**) Percentage of chromatin loop anchors in PP2 containing only class I, only class II, or both class I and class II enhancers. (**E**) Percentage of genes associated with only class I, only class II, or both class I and class II enhancers. Target genes were assigned to enhancers based on nearest TSS of expressed genes (fragments per kilobase per million fragments mapped (FPKM) ≥ 1) in PP2. (**F**) Schematic illustrating FOXA1/2 occupancy, chromatin accessibility, and presence of H3K4me1 and H3K27ac at class I and class II enhancers in GT and PP2. All ChIP-seq and ATAC-seq experiments, n = 2 replicates from independent differentiations. See also Figure S3 and Table S3.

We next sought to determine whether class I and class II enhancers function together within larger regions of active chromatin such as super-enhancers (Whyte et al., 2013), or whether they reside in distinct regulatory domains. To distinguish between these possibilities, we defined 167 super-enhancers among the 2574 pancreatic enhancers identified in Figure S2D (**Figure S3C** and **Table S3A**) and found that 160 (96%) were FOXA1/2-bound at the PP2 stage (**Figure S3D**). Analysis of overlap between class I or class II enhancers and FOXA-bound super-enhancers revealed that most FOXA-bound super-enhancers (76%) contained either class I or class II enhancers but not both (**Figure 3C**). Furthermore, we analyzed Hi-C datasets produced from PP2 stage cells and found that class I and class II enhancers were mostly located in non-overlapping 3D chromatin loops (**Figure 3D** and **Table S3B**). This evidence indicates that class I and class II enhancers reside largely within distinct gene regulatory domains and therefore likely function independently.

To identify target genes of class I and class II enhancers, we assigned enhancers to their nearest expressed gene at the PP2 stage (**Table S3C** and **S3D**), and validated predictions by showing regulation of these genes by FOXA1/2 (**Figure S3E**). Consistent with their location in distinct regulatory domains (**Figures 3C** and **3D**), class I and class II enhancers mostly associated with distinct genes, including pancreatic lineage-determining TFs (**Figure 3E**). Together, these results suggest that two distinct mechanisms establish the pancreatic gene expression program: a subset of pancreatic genes are regulated by enhancers that undergo FOXA1/2-mediated chromatin priming in gut tube, whereas the majority of pancreatic genes are regulated by enhancers that are unprimed prior to pancreatic lineage induction, and to which FOXA1/2 are recruited upon lineage induction (**Figure 3F**).

### Distinct DNA sequence motifs at primed and unprimed pancreatic enhancers

We next investigated mechanisms that could explain the observed temporal differences in FOXA1/2 binding to class I (primed) and class II (unprimed) pancreatic enhancers. To test whether differences in DNA sequence could provide an explanation, we conducted *de novo* motif analysis to identify motifs enriched at class I enhancers against a background of class II enhancers. Class I enhancers were enriched for FOXA motifs and motifs for several signal-dependent TFs, including the ETS family TFs GABPA and SPDEF, the downstream effector of Hippo signaling TEAD, and the retinoic acid receptor RXRA (**Figure 4A** and **Table S4A**). Work in model organisms has identified critical roles for ETS TFs as well as Hippo and retinoic acid signaling in early pancreatic development (Cebola et al., 2015; Huang et al., 2014; Kobberup et al., 2007; Mamidi et al., 2018), suggesting that pancreatic lineage-inductive signals are read at class I enhancers by partnering of FOXA1/2 with signal-dependent TFs.

**Figure 4.**
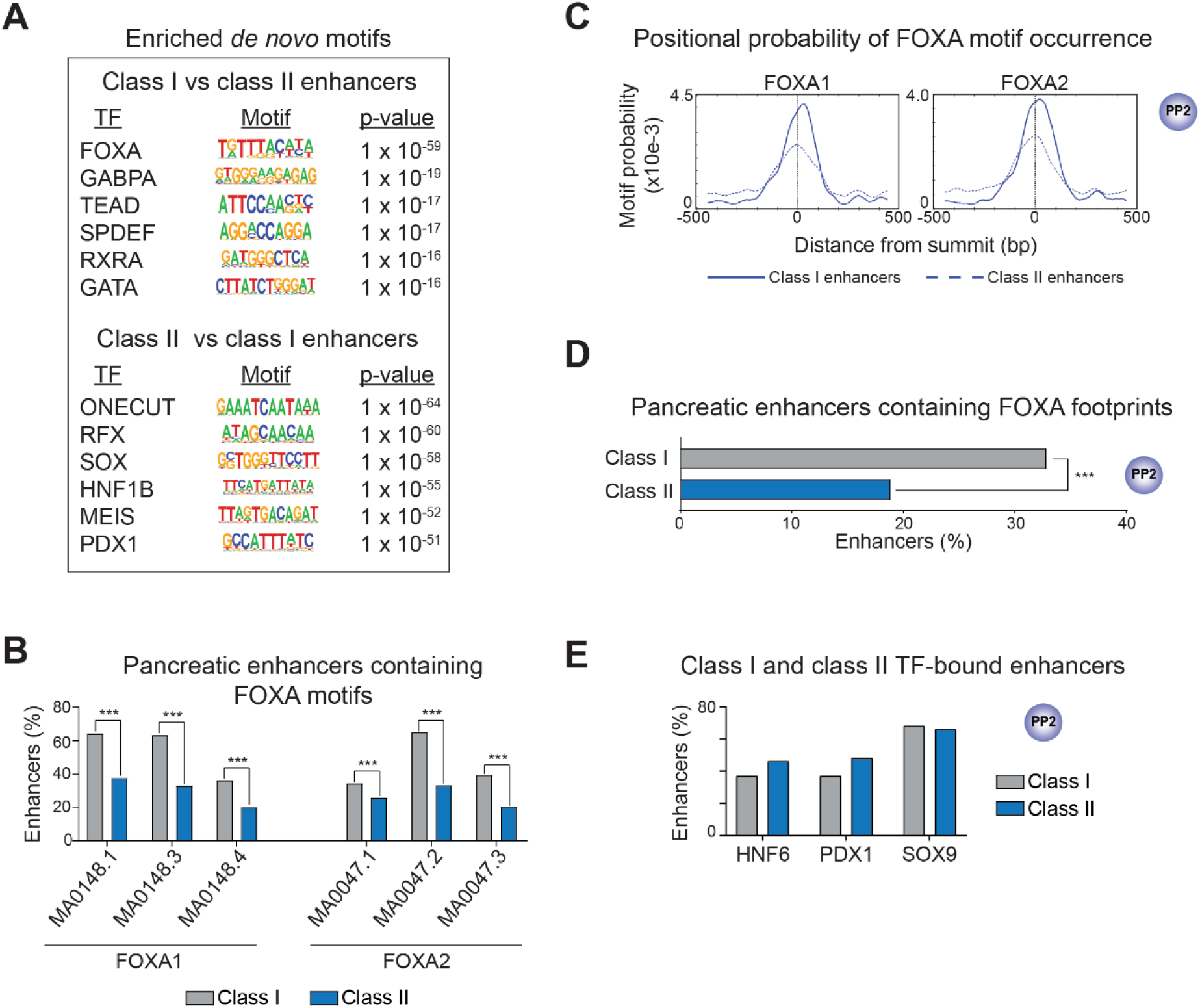
FOXA1/2 binding sites at class I and class II pancreatic enhancers differ in DNA sequence. (**A**) Enriched *de novo* transcription factor (TF) binding motifs at class I against a background of class II pancreatic enhancers and vice versa. Fisher’s exact test, 1-sided, corrected for multiple comparisons. (**B**) Percentage of class I and class II enhancers with at least one occurrence of selected FOXA1 and FOXA2 motifs (*P* = < 2.2 x 10^-16^, < 2.2 x 10^-16^, 1.76 x 10^-13^, 1.61 x 10^-4^, < 2.2 x 10^-16^, and < 2.2 x 10^-16^ for comparisons of occurrences of MA0148.1, MA0148.3, MA0148.4, MA0047.1, MA0047.2, and MA0047.3, respectively. Fisher’s exact test, 2-sided). (**C**) Probability (motif occurrence per base pair) of FOXA1 (MA0148.3) and FOXA2 (MA0047.2) motifs relative to ATAC-seq peak summits at class I (solid line) and class II (dashed line) enhancers. ATAC-seq peak summits at class I enhancers are enriched for occurrences of MA0148.3 (*P* = 2.1 x 10^-14^; Fisher’s exact test, 1-sided) and MA0047.2 (*P* = 6.8 x 10^-14^) compared to summits at class II enhancers. (**D**) Percentage of class I and class II enhancers containing FOXA TF ATAC-seq footprints in PP2 (*P* = 1.01 x 10^-10^ for comparison of class I and class II enhancers; Fisher’s exact test, 2-sided). (**E**) Percentage of class I and class II enhancers overlapping HNF6, PDX1, and SOX9 ChIP-seq peaks (within 100 bp from peak) in PP2. All ChIP-seq experiments, n = 2 replicates from independent differentiations. See also Figure S4 and Table S4.

Since FOXA1/2 binding to class I enhancers precedes binding to class II enhancers (**Figure 2A**) and FOXA motifs are enriched at class I compared to class II enhancers (**Figure 4A**), we postulated that different mechanisms could underlie FOXA1/2 recruitment to the two classes of enhancers. Given that binding site selection of pioneer TFs such as FOXA1/2 has been shown to depend on motif abundance, strength, and position (Donaghey et al., 2018; Farley et al., 2015; Swinstead et al., 2016; Zhang et al., 2015), we analyzed FOXA motifs at class I and class II enhancers for these features. To determine abundance and strength of FOXA motifs, we selected position-weighted matrices (PWMs) corresponding to three FOXA1 and three FOXA2 motifs from JASPAR (Fornes et al., 2020) (**Figure S4A**), identified occurrences of each motif at class I and class II enhancers, and generated a log-odds score to measure how closely the DNA sequence at each identified motif occurrence matched the PWM. Class I enhancers were significantly enriched for occurrences of all six FOXA motifs compared to class II enhancers (**Figure 4B**). Furthermore, three of the FOXA motifs had significantly higher log-odds scores at class I than class II enhancer occurrences (MA0047.2, MA0148.1, and MA0148.3; *P* = 1.54 x 10^-2^, 1.10 x 10^-3^, and 1.03 x 10^-2^, respectively; Wilcoxon rank sum test). Thus, class II enhancers contain more degenerate and fewer FOXA motifs compared to class I enhancers. We additionally examined the positioning of FOXA motifs relative to open chromatin by identifying regions of greatest chromatin accessibility at class I and class II enhancers in PP2 stage cells (n = 531 and n = 1257 ATAC-seq summits in class I and class II enhancers, respectively) and determining enrichment of each FOXA motif at these regions. Occurrence of all FOXA motifs was enriched at ATAC-seq summits at class I compared to class II enhancers (**Figures 4C** and **S4B**), indicating that regions of greatest chromatin accessibility at class I enhancers are more likely to harbor FOXA motifs. ATAC-seq footprinting analysis further revealed a higher occurrence of FOXA footprints at class I than at class II enhancers (**Figure 4D**), indicative of either longer FOXA1/2 DNA residence times or more direct interaction of FOXA1/2 with DNA at class I enhancers (Sung et al., 2014). Together, this analysis reveals features of FOXA motifs at class I pancreatic enhancers previously associated with canonical FOXA1/2 pioneer TF activity (Donaghey et al., 2018; Swinstead et al., 2016).

To further elucidate differences in mechanisms of FOXA recruitment to class I and class II enhancers, we identified *de novo* motifs enriched at class II enhancers against a background of class I enhancers. Here, we observed enrichment of motifs for pancreatic lineage-determining TFs, such as ONECUT (HNF6), SOX (SOX9), HNF1B, and PDX1 (**Figure 4A** and **Table S4B**), which sharply increased in expression during pancreatic lineage induction (**Figure S4C**). To determine whether these TFs exhibit preferential binding to class II enhancers, as implied by differences in motif enrichment, we mapped HNF6, PDX1, and SOX9 binding sites genome-wide at the PP2 stage (**Figures 4E** and **S4D**). Surprisingly, similar percentages of class I and class II enhancers were bound by HNF6, PDX1, and SOX9 (**Figure 4E**). To determine whether the difference in sequence motif enrichment between class I and class II enhancers is also observed when focusing on enhancers bound by a specific TF, we analyzed motifs at HNF6-, PDX1-, or SOX9- bound enhancers. Still, class I enhancers were enriched for FOXA and class II enhancers for ONECUT (HNF6), PDX1, and SOX motifs (**Figure S4E** and **Table S4C-S4H**). Thus, despite differences in DNA sequence motifs between primed (class I) and unprimed (class II) enhancers, both classes of enhancers are occupied by FOXA1/2 as well as pancreatic lineage-determining TFs after pancreatic lineage induction.

### FOXA1/2 binding to unprimed but not primed enhancers depends on PDX1

Since motifs for pancreatic lineage-determining TFs, such as PDX1, were enriched at class II compared to class I enhancers (**Figure 4A**), we hypothesized that FOXA1/2 recruitment to class II enhancers could require cooperativity with lineage-determining TFs. To test this, we analyzed FOXA1/2 binding, chromatin accessibility, and H3K27ac signal in *PDX1*-deficient pancreatic progenitors (**Figures 5A** and **S5A**). Focusing on PDX1-bound enhancers (n = 205 class I enhancers and 682 class II enhancers), we found that loss of *PDX1* reduced FOXA1/2 binding to a greater extent at class II than class I enhancers (**Figure 5B**), exemplified by class I enhancers near *PDX1* and *HNF1B*, and class II enhancers near *NKX6.1* and *MNX1* (**Figure 5C**). In total, 92% of class I enhancers compared to 68% of class II enhancers maintained a FOXA1/2 ChIP-seq peak after *PDX1* knock-down (**Figure S5B**). Given substantial overlap between binding sites for pancreatic lineage-determining TFs (**Figure S4D**), it is possible that other TFs recruit FOXA1/2 at PDX1-bound class II enhancers where FOXA1/2 occupancy is maintained. Loss of *PDX1* led to a significant reduction in ATAC-seq and H3K27ac signal at both class I and class II enhancers (**Figure S5C**), showing that full acquisition of chromatin accessibility and enhancer activation during pancreas induction require PDX1 at primed and unprimed enhancers.

**Figure 5.**
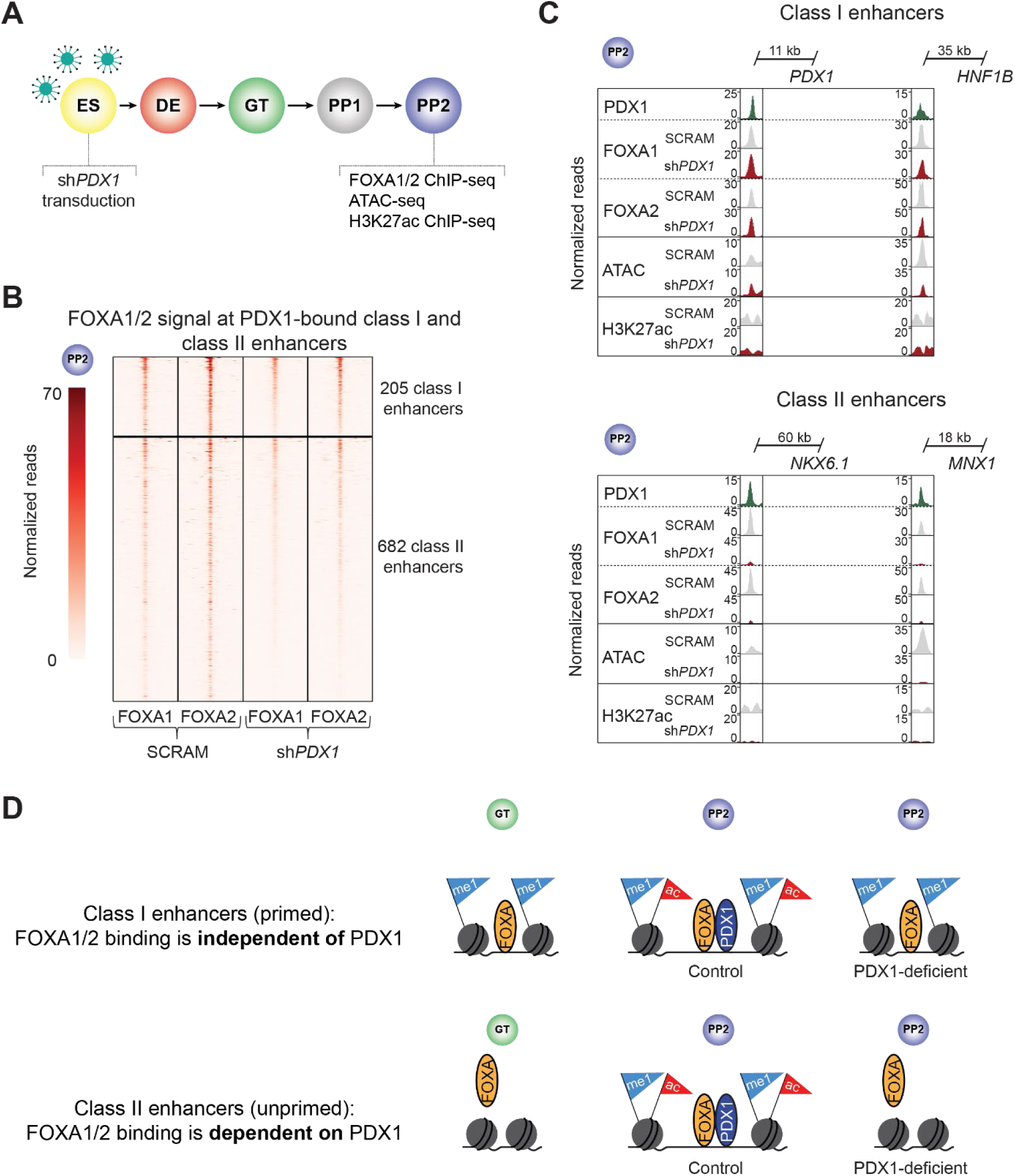
FOXA1/2 binding at class II enhancers is dependent on PDX1. (**A**) Schematic of experimental design for *PDX1* knock-down in hESCs and subsequent differentiation into PP2 stage pancreatic progenitors. (**B**) Heatmap showing density of FOXA1 and FOXA2 ChIP-seq reads at PDX1-bound class I and class II pancreatic enhancers in hESCs transduced with scrambled control (SCRAM) or *PDX1* shRNA (sh*PDX1*) in PP2. Heatmap is centered on FOXA1 and FOXA2 peaks, respectively, and spans 5 kb. (**C**) Genome browser snapshots showing PDX1, FOXA1, and FOXA2 ChIP-seq, ATAC-seq, and H3K27ac ChIP-seq signal at class I enhancers near *PDX1* and *HNF1B* and class II enhancers near *NKX6.1* and *MNX1* in PP2. Approximate distance between enhancer and gene body is indicated. (**D**) Schematic illustrating distinct modes of FOXA TF recruitment at class I and class II pancreatic enhancers. FOXA1/2 recruitment depends on the lineage-determining TF PDX1 at class II enhancers. Both enhancer classes require PDX1 or activation. All ChIP-seq and ATAC-seq experiments, n = 2 replicates from independent differentiations. See also Figure S5.

Collectively, our findings show that despite similar mechanisms for their activation, primed and unprimed pancreatic enhancers differ in sequence logic and mechanism of FOXA1/2 recruitment (**Figure 5D**). Primed enhancers have abundant and strong FOXA motifs, and FOXA1/2 are recruited to primed enhancers prior to pancreatic lineage induction independent of the pancreatic TF PDX1. By contrast, unprimed enhancers have fewer and weaker FOXA motifs, and FOXA1/2 are recruited to unprimed enhancers by PDX1 concomitant with lineage induction.

### Distinct temporal patterns of FOXA1/2 occupancy distinguish hepatic and alveolar enhancers

To determine whether the identified mechanisms of enhancer activation during organ development are universal across endodermal lineages, we also analyzed liver and lung enhancers, which like pancreatic enhancers undergo chromatin priming in gut endoderm (Wang et al., 2015). Similar to pancreas development, both early liver and lung development depend on FOXA TFs (Genga et al., 2019; Lee et al., 2005; Wan et al., 2005). Furthermore, previous studies have demonstrated FOXA binding to primed liver enhancers in gut endoderm prior to organ lineage induction (Gualdi et al., 1996; Wang et al., 2015). To test whether class I and class II enhancers can be distinguished during liver and lung development, we induced the hepatic fate from hESC-GT stage intermediates (**Figure 6A**), and generated distal lung alveolar epithelial type 2-like cells (iAT2s) grown at 95% purity as 3D alveolospheres (ALV) from iPSCs (**Figure 6B**) (Jacob et al., 2017; Wang et al., 2015). For liver, we analyzed H3K27ac signal and FOXA1/2 binding before liver induction at the GT stage and in hepatic progenitors (HP). For lung, we analyzed H3K27ac signal and FOXA1 binding before lung induction in anteriorized foregut (AFG) and at the ALV stage.

**Figure 6.**
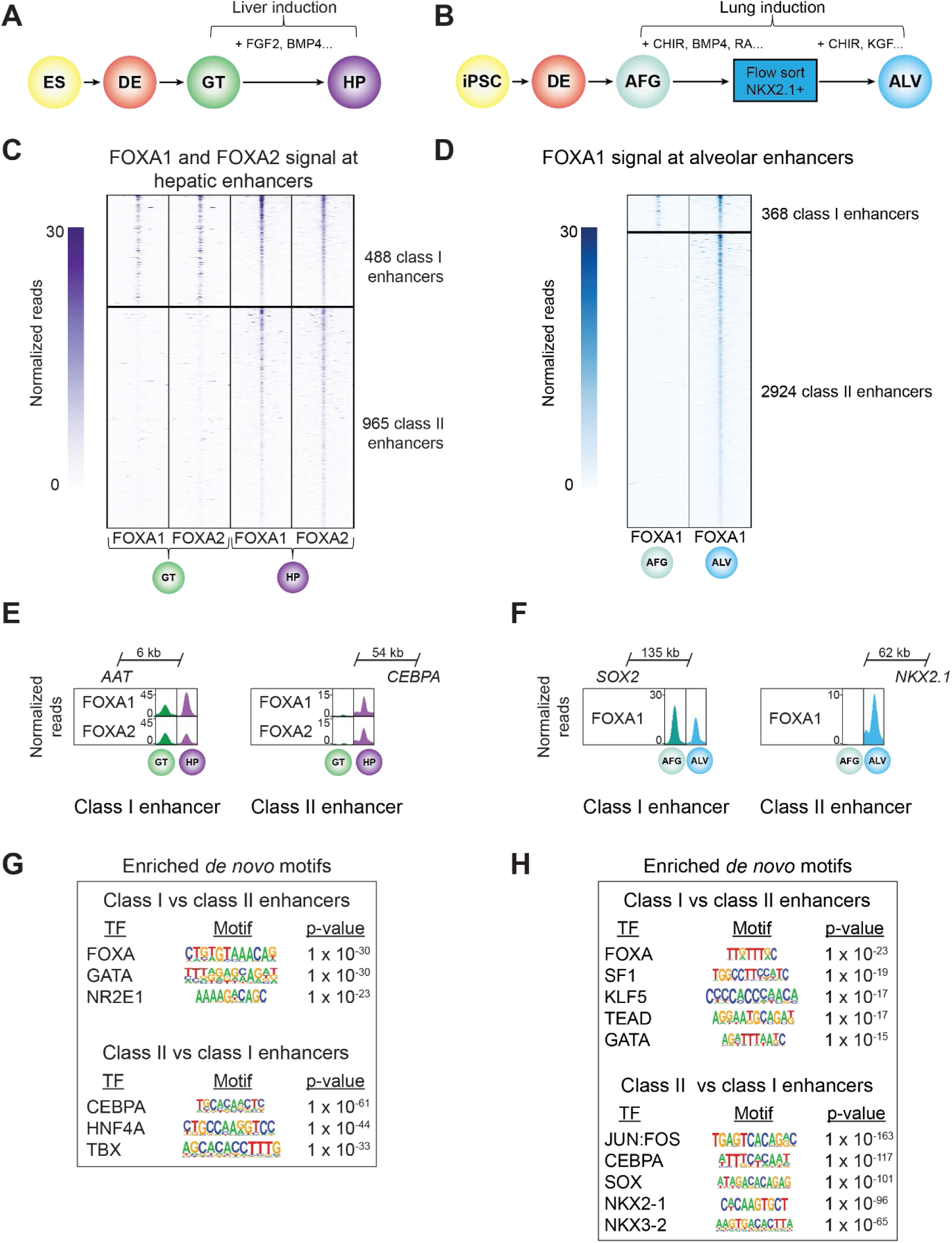
Class I and class II enhancers can be distinguished in liver and lung development. (**A** and **B)** Schematic of stepwise differentiation of hESCs to hepatic progenitors (HP) (**A**) and induced human pluripotent stem cells (iPSC) into alveolosphere organoids (ALV) (**B**). AFG, anteriorized foregut. Select growth factors for hepatic (**A**) and alveolar (**B**) lineage induction are indicated. FGF2, fibroblast growth factor 2; BMP4, bone morphogenic protein 4; CHIR, CHIR99021; RA, retinoic acid. (**C**) Heatmap showing density of FOXA1 and FOXA2 ChIP-seq reads at hepatic enhancers in GT and HP. Heatmap is centered on FOXA1 and FOXA2 peaks, respectively, and spans 5 kb. Hepatic enhancers are classified based on temporal pattern of FOXA1 and FOXA2 occupancy. (**D**) Heatmap showing density of FOXA1 ChIP-seq reads at hepatic enhancers in AFG and ALV. Heatmap is centered on FOXA1 peaks and spans 5 kb. Alveolar enhancers are classified based on temporal pattern of FOXA1 occupancy. (**E**) Genome browser snapshots showing FOXA1 and FOXA2 ChIP-seq signal at a class I hepatic enhancer near *AAT* and a class II hepatic enhancers near *CEBPA* in GT and HP. (**F**) Genome browser snapshots showing FOXA1 ChIP-seq signal at a class I alveolar enhancer near *SOX2* and a class II alveolar enhancer near *NKX2.1* in AFG and ALV. (**G** and **H**) Enriched *de novo* transcription factor (TF) binding motifs at class I against a background of class II enhancers and vice versa for hepatic (**G**) and alveolar enhancers (**H**). Fisher’s exact test, 1-sided, corrected for multiple comparisons. All ChIP-seq experiments, n = 2 replicates from independent differentiations. See also Figures S6 and S7, Tables S5 and S6.

Analogous to the strategy used for identifying pancreatic enhancers (**Figure S2D**), we identified hepatic and alveolar enhancers based on gain in H3K27ac signal during the GT to HP and AFG to ALV transitions, respectively (≥ 2 fold change in H3K27ac, FDR ≤ 0.05; **Figures S6A-S6D**). Subsequently, we quantified FOXA1/2 binding at the identified enhancers. As in pancreas, we observed two distinct patterns of FOXA1/2 occupancy (**Figures 6C, 6D** and **Table S5**) despite similar dynamics in H3K27ac signal (**Figures S6E** and **S6F**): a subset of class I enhancers exhibited FOXA1/2 occupancy prior to lineage induction (488 class I hepatic enhancers and 368 class I alveolar enhancers), whereas class II enhancers constituted the majority and exhibited *de novo* FOXA1/2 binding with lineage induction (965 class II hepatic enhancers and 2924 class II alveolar enhancers). These patterns were exemplified by enhancers near hepatic genes Alpha1-Antitrypsin (*AAT*) and *CEBPA* (**Figure 6E**), as well as lung developmental TF genes *SOX2* and *NKX2.1* (**Figure 6F**).

*De novo* motif analysis at class I against a background of class II hepatic enhancers revealed enrichment for FOXA motifs and the motif for the signal-dependent nuclear receptor NR2E1 (Corso-Diaz et al., 2016). Class II enhancers showed comparative enrichment for motifs of the hepatic lineage-determining TFs CEBPA, HNF4A, and TBX (Kheolamai and Dickson, 2009; Papaioannou, 2014) (**Figure 6G**, **Tables S6A** and **S6B**), which increased in expression upon liver induction from hESC-GT intermediates (**Figure S7A**). FOXA2, HNF4A, and CEBP have been shown to co-bind liver-specific enhancers after liver induction (Iwafuchi-Doi et al., 2016), supporting a potential role for cooperative recruitment of FOXA TFs by these factors. Analogous to the motif enrichment patterns observed in pancreas and liver, alveolar class I enhancers were comparatively enriched for FOXA motifs and motifs for signal-dependent TFs NR5A1 (SF1) and TEAD with roles in lung development (Ikonomou et al., 2020; Ramana, 2019), whereas alveolar class II enhancers showed comparative motif enrichment for SOX family TFs and the lung master TF NKX2.1 (Herriges and Morrisey, 2014) (**Figure 6H**, **Tables S6C** and **S6D**). Thus, as in pancreas, a subset of hepatic and alveolar enhancers with canonical FOXA motifs and enrichment for motifs of signal-dependent TFs are FOXA1/2- bound prior to lineage induction, while *de novo* FOXA1/2 recruitment occurs at the majority of hepatic and alveolar enhancers upon lineage induction.

To gain further insight into the architecture of hepatic and alveolar enhancers, we examined abundance, strength, and positioning of FOXA motifs. Using the same six FOXA PWMs as for pancreatic enhancers (**Figure S4A**), we observed significant enrichment for occurrence of FOXA motifs at both class I hepatic and class I alveolar enhancers (**Figures S7B** and **S7C**). We also found significantly higher log-odds scores for three FOXA PWMs (MA0047.2, MA0148.1, and MA0148.3; *P* = 1.40 x 10^-3^, 2.00 x 10^-3^, and 1.60 x 10^-2^, respectively; Wilcoxon rank sum test) at class I compared to class II hepatic enhancers, and two FOXA PWMs (MA0047.3 and MA0148.1; *P* = 3.1 x 10^-2^ and 4.1 x 10^-2^, respectively; Wilcoxon rank sum test) at class I compared to class II alveolar enhancers. Furthermore, FOXA motif occurrence at ATAC-seq summits (444 and 701 ATAC-seq summits in class I and class II enhancers, respectively, at HP stage; **Figure S7D**) and occurrence of FOXA footprints (**Figure S7E**) were enriched at class I compared to class II hepatic enhancers. Thus, similar to pancreatic class I enhancers, hepatic,, and alveolar class I enhancers exhibit sequence features that have been associated with canonical FOXA1/2 pioneering in other contexts (Donaghey et al., 2018; Swinstead et al., 2016). Moreover, analogous to pancreatic class II enhancers, we observed no binding preference of the hepatic lineage-determining TF HNF4A at class II compared to class I hepatic enhancers despite HNF4A motif enrichment at HNF4A-bound class II enhancers (**Figures S7F**, **S7G**, **Tables S6E** and **S6F**). These results show that similar characteristics of sequence architecture distinguish pancreatic, hepatic, and alveolar class I and class II enhancers. Across all three lineages, class I enhancers harbor more abundant and stronger FOXA motifs and are enriched for motifs of signal-dependent TFs, while class II enhancers are enriched for motifs of lineage-determining TFs.

### Lineage-specific recruitment of FOXA1/2 to unprimed enhancers

Our results suggest a model whereby the full enhancer complement for each endodermal organ lineage is established through (i) FOXA1/2-mediated priming of a small subset of enhancers for each lineage in endodermal precursors prior to lineage induction, and (ii) activation of a larger subset of unprimed enhancers by organ lineage-determining TFs that cooperatively recruit FOXA1/2 upon lineage induction. To determine the relationship between class I and class II enhancers across different endodermal lineages, we performed differential motif enrichment analysis, comparing class I enhancers or class II enhancers of each lineage against a background of class I enhancers or class II enhancers, respectively, of the alternate lineages. As expected, motifs for lineage-determining TFs for each lineage were enriched at both classes of enhancers (**Tables S7A-F**). However, motif enrichment was stronger at class II than at class I enhancers (**Figure 7A**), lending further support to the model that cooperativity with lineage-determining TFs facilitates lineage-specific FOXA1/2 association with class II enhancers of each organ. Consistent with the binding of FOXA to class I enhancers in shared developmental precursors prior to lineage induction, we found that class I enhancers of one organ lineage were more frequently bound by FOXA1/2 in alternate lineages than class II enhancers (**Figures 7B** and **7C**). Altogether, these findings support establishment of organ-specific gene expression programs through two distinct mechanisms of FOXA1/2-mediated enhancer activation (**Figure 7D**).

**Figure 7.**
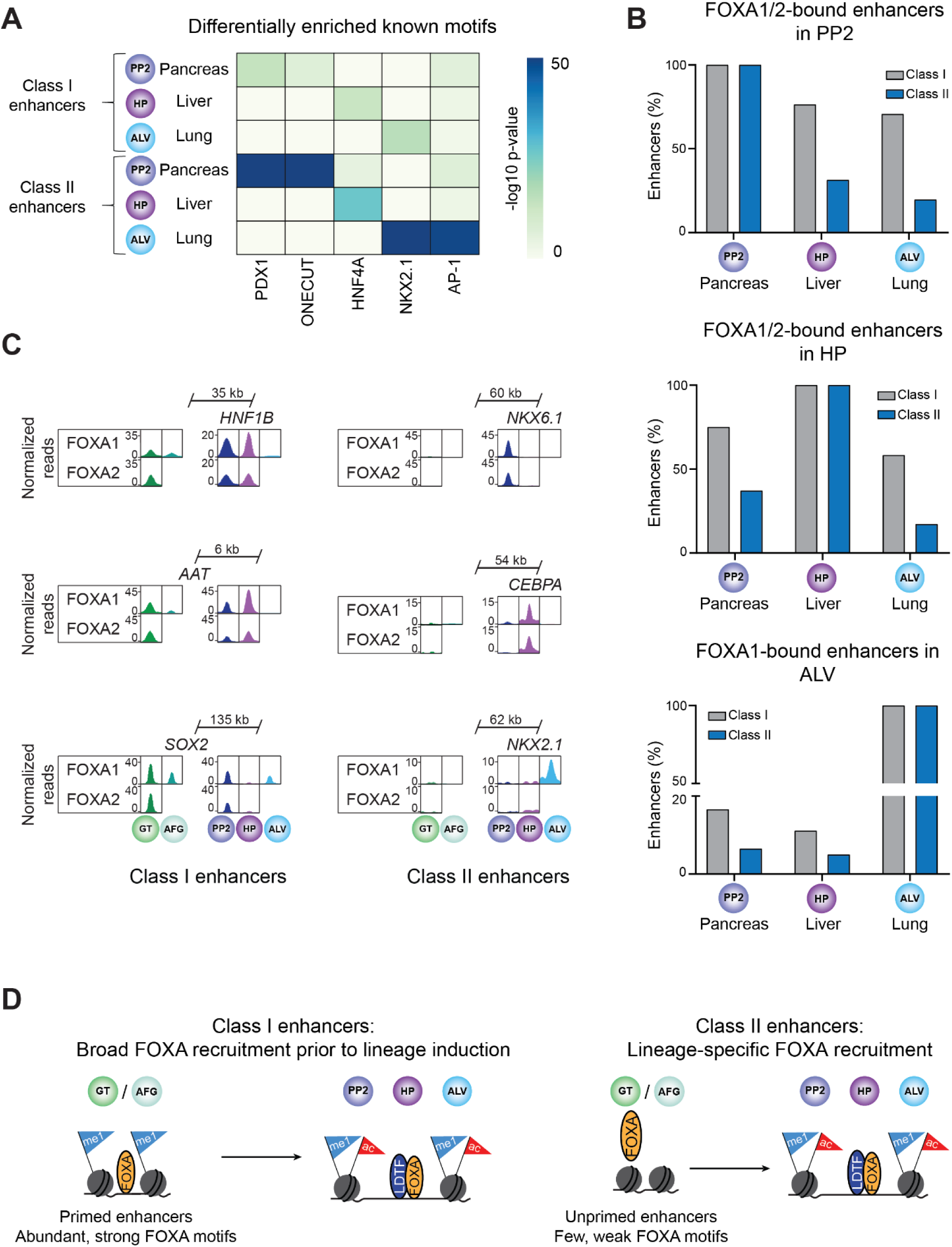
Recruitment of FOXA1/2 to class II enhancers is lineage-specific. (**A**) Heatmap showing enrichment of known binding motifs for lineage-determining transcription factors at pancreatic, hepatic, and alveolar class I and class II enhancers. Class I and class II enhancers of each lineage were compared against a background of class I and class II enhancers, respectively, of all other lineages. Fisher’s exact test, 1-sided, corrected for multiple comparisons. (**B**) Percentage of pancreatic, hepatic, and alveolar class I and class II enhancers overlapping FOXA1/2 ChIP-seq peaks (within 100 bp from peak) in PP2 (pancreas), HP (liver) and ALV (lung). For ALV only FOXA1 peaks were considered. (**C**) Genome browser snapshots showing FOXA1/2 ChIP-seq signal across endodermal lineages at example pancreatic, hepatic, and alveolar class I and class II enhancers. Approximate distance between enhancer and gene body is indicated. (**D**) Schematic showing differential recruitment of FOXA TFs to endodermal organ class I and class II enhancers during endoderm development. LDTF, lineage determining transcription factors. All ChIP-seq experiments, n = 2 replicates from independent differentiations. See also Table S7.

## DISCUSSION

Chromatin priming at enhancers is broadly observed in development and is defined by chromatin remodeling at lineage-specific enhancers prior to enhancer activation and target gene expression (Bonifer and Cockerill, 2017; Creyghton et al., 2010; Rada-Iglesias et al., 2011). We have previously reported that chromatin priming and FOXA1/2 recruitment precedes organ lineage induction at pancreas, liver, and lung enhancers (Wang et al., 2015). Here, we show that chromatin priming and early FOXA1/2 recruitment is limited to a small subset of organ lineage enhancers, whereas the majority transition from unprimed to active and engage FOXA1/2 upon lineage induction. We demonstrate that primed enhancers can be distinguished from unprimed enhancers based on sequence logic and mechanism of FOXA1/2 recruitment. The results presented here provide a molecular framework for understanding how different organ-specific gene expression programs are established from a common developmental precursor.

It has been shown that the ability of FOXA2 to stably bind and remodel chromatin is DNA sequence-dependent. At FOXA2 binding sites where ectopic FOXA2 expression can induce DNA accessibility, FOXA motifs are specifically enriched and widely distributed across the binding region (Donaghey et al., 2018). Conversely, FOXA1 binding to sites poorly enriched for canonical FOXA1 motifs in breast cancer cell lines requires steroid receptor activation (Paakinaho et al., 2019; Swinstead et al., 2016), suggesting that FOXA1 recruitment to sites with low abundance of FOXA1 motifs depends on additional TFs. Consistent with these findings, we observed stronger and more abundant FOXA motifs at primed compared to unprimed enhancers and found that FOXA1/2 recruitment to unprimed pancreatic enhancers depends on the pancreatic lineage-determining TF PDX1 which becomes expressed upon pancreatic lineage induction. Thus, our results extend previous studies in immortalized cell lines to show that distinct sequence architecture and mechanisms of FOXA recruitment exist within an organ-specific enhancer repertoire, and that enhancers harboring different sequence features can be distinguished based on temporal patterns of developmental FOXA recruitment. The temporal differences in FOXA1/2 occupancy raise the question of whether primed and unprimed enhancers control different gene regulatory programs. Although we found primed and unprimed enhancers to associate to a large extent with different genes, we observed no clear difference in the function or expression of genes predicted to be regulated by the two enhancer classes. Distinct early pancreatic TFs, with comparable temporal expression patterns, associate with primed (e.g. *HNF1B* and *GLIS3*) or unprimed enhancers (e.g. *SOX9* and *MNX1*). Thus, consistent with the similar activity pattern of primed and unprimed enhancers, there does not appear to be a temporal difference in the expression of their target genes.

*In vitro* assays have shown that FOXA1/2 can engage target sites on nucleosomal DNA, which is referred to as pioneering activity (Cirillo et al., 2002; Iwafuchi-Doi et al., 2016; Soufi et al., 2015). The importance of FOXA TF pioneering activity for target site engagement in cellular contexts is the subject of an ongoing debate (Iwafuchi et al., 2020; Iwafuchi-Doi et al., 2016; Paakinaho et al., 2019; Soufi et al., 2015; Swinstead et al., 2016). The higher abundance of FOXA footprints at primed enhancers is consistent with longer residence times of FOXA TFs, a feature that has been associated with pioneering activity (Sekiya et al., 2009). However, cooperative binding with GATA TFs could also play a role, as GATA motifs were enriched at primed compared to unprimed pancreatic, hepatic, and alveolar enhancers (**Figures 4A, 6G** and **6H**), and GATA4 is known to facilitate FOXA2 binding in other contexts (Donaghey et al., 2018). Of note, pioneering activity and cooperative binding are not mutually exclusive events (Iwafuchi et al., 2020). Regardless of the precise mechanism by which FOXA1/2 contact their target sites, the enrichment of motifs for organ lineage-determining TFs at unprimed enhancers suggests relevance of lineage-specific TFs for FOXA1/2 binding to these enhancers across endodermal organs, as shown for PDX1 at pancreatic enhancers.

To initiate organ-specific gene expression during development, a subset of enhancers must be able to respond to environmental cues that induce the organ fate. The motif enrichment for signal-dependent TFs at primed enhancers suggests that lineage-inductive cues are read at the level of primed enhancers, agreeing with prior observations that chromatin priming at pancreatic, hepatic, and lung enhancers coincides with the acquisition of responsiveness to lineage-inductive cues (Wang et al., 2015). Given that priming of enhancers occurs broadly across endodermal organ lineages, differences in organ-inductive signals could contribute to specificity of enhancer activation. While enhancer priming may facilitate enhancer activation upon exposure to lineage-inductive cues, the persistence of the primed state at enhancers for alternative fates after lineage induction (**Figure 7B**) could also bear a risk for aberrant gene activation by subsequent developmental exposure to signals or TFs able to activate these enhancers. Conversely, the indirect recruitment of FOXA by lineage-specific TFs to the majority of organ-specific enhancers at sites with suboptimal FOXA motifs could provide a safeguard against ectopic gene activation after lineage commitment. In agreement with this concept, FOXA2 peaks at liver-specific genes are associated with weaker FOXA2 motifs than FOXA2 peaks at broadly expressed genes (Tuteja et al., 2008). Furthermore, studies in *Drosophila* and *Ciona* suggest that suboptimization of TF binding motifs could be a general principle by which to confer cell specificity to enhancers (Crocker et al., 2015; Farley et al., 2015).

Previous studies have shown that as cells adopt a particular fate, they lose the ability to respond to inductive cues for alternate fates (Rankin et al., 2018; Wang et al., 2015; Zorn and Wells, 2009). In this study, we observed loss of FOXA1/2 binding at 24% of primed hepatic enhancers upon pancreas induction, and at 25% of primed pancreatic enhancers upon liver induction. Thus, our data suggest that commitment to a given lineage is accompanied by the eviction of FOXA1/2 from at least a subset of enhancers associated with alternative lineages. The extent to which FOXA1/2 are further evicted as organ-committed cells terminally differentiate and mature remains to be determined. Exploring a possible link between residual primed enhancers for alternative lineages and cell plasticity is an interesting direction for future studies. For example, cells of related developmental origin can interconvert to regenerate lost tissue cells (Deng et al., 2018), and oncogenic transformation of endodermal organs is associated with the activation of genes normally expressed in developmentally related organs (Tata et al., 2018). Understanding the mechanisms that confer and restrict the ability to activate lineage-specific enhancers will not only help generate accurate hPSC models, but also provide insights into mechanisms of gene regulation in disease.

## METHODS

### CONTACT FOR REAGENT AND RESOURCE SHARING

Further information and requests for resources and reagents should be directed to and will be fulfilled by the Lead Contact, Maike Sander.

### EXPERIMENTAL MODEL AND SUBJECT DETAILS

#### Human cell culture experiments

hESC research was approved by the University of California, San Diego (UCSD), Institutional Review Board and Embryonic Stem Cell Research Oversight Committee (protocol 090165ZX). Human iPSC research was approved by the Boston University Institutional Review Board (protocol H-33122).

#### Maintenance of HEK293T cells

HEK293T cells (female) were cultured in a humidified incubator at 37 °C with 5% CO_2_ using Dulbecco’s Modified Eagle Medium (Cat# 45000-312; 4.5 g/L glucose, [+] L-glutamine, [-] sodium pyruvate) supplemented with 10% fetal bovine serum (FBS).

#### Maintenance and differentiation of CyT49 hESCs

CyT49 hESCs (male) were maintained and differentiated as described (Schulz et al., 2012; Wang et al., 2015; Xie et al., 2013). Propagation of CyT49 hESCs was carried out by passing cells every 3 to 4 days using Accutase™ (eBioscience) for enzymatic cell dissociation, and with 10% (v/v) human AB serum (Valley Biomedical) included in the hESC media the day of passage. hESCs were seeded into tissue culture flasks at a density of 50,000 cells/cm^2^.

Pancreatic differentiation was performed as previously described (Schulz et al., 2012; Wang et al., 2015; Xie et al., 2013). Briefly, a suspension-based culture format was used to differentiate cells in aggregate form. Undifferentiated aggregates of hESCs were formed by re-suspending dissociated cells in hESC maintenance medium at a concentration of 1 x 10^6^ cells/mL and plating 5.5 mL per well of the cell suspension in 6- well ultra-low attachment plates (Costar). The cells were cultured overnight on an orbital rotator (Innova2000, New Brunswick Scientific) at 95 rpm. After 24 hours the undifferentiated aggregates were washed once with RPMI medium and supplied with 5.5 mL of day 0 differentiation medium. Thereafter, cells were supplied with the fresh medium for the appropriate day of differentiation (see below). Cells were continually rotated at 95 rpm, or 105 rpm on days 4 through 8, and no media change was performed on day 10. Both RPMI (Mediatech) and DMEM High Glucose (HyClone) medium were supplemented with 1X GlutaMAX™ and 1% penicillin/streptomycin. Human activin A, mouse Wnt3a, human KGF, human noggin, and human EGF were purchased from R&D systems. Other added components included FBS (HyClone), B-27® supplement (Life Technologies), Insulin-Transferrin-Selenium (ITS; Life Technologies), TGFβ R1 kinase inhibitor IV (EMD Bioscience), KAAD-Cyclopamine (KC; Toronto Research Chemicals), and the retinoic receptor agonist TTNPB (RA; Sigma Aldrich). Day-specific differentiation media formulations were as follows:

Days 0 and 1: RPMI + 0.2% (v/v) FBS, 100 ng/mL Activin, 50 ng/mL mouse Wnt3a, 1:5000 ITS. Days 1 and 2: RPMI + 0.2% (v/v) FBS, 100 ng/mL Activin, 1:5000 ITS
Days 2 and 3: RPMI + 0.2% (v/v) FBS, 2.5 mM TGFβ R1 kinase inhibitor IV, 25ng/mL KGF, 1:1000 ITS
Days 3 - 5: RPMI + 0.2% (v/v) FBS, 25 ng/mL KGF, 1:1000 ITS
Days 5 - 8: DMEM + 0.5X B-27® Supplement, 3 nM TTNPB, 0.25 mM KAAD-Cyclopamine, 50 ng/mL Noggin
Days 8 - 10: DMEM/B-27, 50 ng/mL KGF, 50 ng/mL EGF

Cells at D0 correspond to the embryonic stem cell (ES) stage, cells at D2 correspond to the definitive endoderm (DE) stage, cells at D5 correspond to the gut tube (GT) stage, cells at D7 correspond to the early pancreatic progenitor (PP1) stage, and cells at D10 correspond to the late pancreatic progenitor (PP2) stage.

Hepatic differentiation was performed as previously described (Wang et al., 2015). Briefly, cells were treated identically as in pancreatic differentiation until the GT stage at D5. At this point cells were treated with 50 ng/ml BMP4 (Millipore) and 10 ng/ml FGF2 (Millipore) in RPMI media (Mediatech) supplemented with 0.2% (vol/vol) FBS (HyClone) for 3 days with daily media changes. Cells at D8 correspond to the hepatic progenitor (HP) cell stage.

#### Maintenance and differentiation of H1 hESCs

H1 hESCs (male) were maintained and differentiated as described with some modifications (Jin et al., 2019; Rezania et al., 2014). In brief, hESCs were cultured in mTeSR1 media (Stem Cell Technologies) and propagated by passaging cells every 3 to 4 days using Accutase (eBioscience) for enzymatic cell dissociation.

For differentiation, cells were dissociated using Accutase for 10 min, then reaggregated by plating the cells at a concentration of ∼5.5 10^6^ cells/well in a low attachment 6-well plate on an orbital shaker (100 rpm) in a 37 °C incubator. The following day, undifferentiated cells were washed in base media (see below) and then differentiated using a multi-step protocol with stage-specific media and daily media changes.

All stage-specific base media were comprised of MCDB 131 medium (Thermo Fisher Scientific) supplemented with NaHCO3, GlutaMAX, D-Glucose, and BSA using the following concentrations:

Stage 1/2 base medium: MCDB 131 medium, 1.5 g/L NaHCO3, 1X GlutaMAX, 10 mM D-Glucose, 0.5% BSA
Stage 3/4 base medium: MCDB 131 medium, 2.5 g/L NaHCO3, 1X GlutaMAX, 10 mM D-glucose, 2% BSA

Media compositions for each stage were as follows:

Stage 1 (days 0 - 2): base medium, 100 ng/ml Activin A, 25 ng/ml Wnt3a (day 0). Day 1- 2: base medium, 100 ng/ml Activin A
Stage 2 (days 3 - 5): base medium, 0.25 mM L-Ascorbic Acid (Vitamin C), 50 ng/mL FGF7
Stage 3 (days 6 - 7): base medium, 0.25 mM L-Ascorbic Acid, 50 ng/mL FGF7, 0.25 µM SANT-1, 1 µM Retinoic Acid, 100 nM LDN193189, 1:200 ITS-X, 200 nM TPB
Stage 4 (days 8 - 10): base medium, 0.25 mM L-Ascorbic Acid, 2 ng/mL FGF7, 0.25 µM SANT-1, 0.1 µM Retinoic Acid, 200 nM LDN193189, 1:200 ITS-X, 100nM TPB

Cells at D0, D3, D6, D8, and D11 correspond to the ES DE, GT, PP1, and PP2 stages, respectively.

#### Maintenance and differentiation of SPC2 iPSCs

SPC2 iPSCs (male; clone SPC2-ST-B2, (Hurley et al., 2020) were maintained in feeder-free culture conditions in 6-well tissue culture dishes (Corning) coated with growth factor reduced Matrigel (Corning), in mTeSR1 medium (Stem Cell Technologies) and passaged using gentle cell dissociation reagent (GCDR). Details of iPSC derivation, characterization, and differentiation into anterior foregut endoderm and alveolar epithelial type 2 cells (iAT2s; also known as iAEC2s) have been previously published (Hurley et al., 2020; Jacob et al., 2017; Jacob et al., 2019) and are available for free download at http://www.bu.edu/dbin/stemcells/protocols.php. Briefly, the SPC2-ST-B2 iPSC clone, engineered to carry a tdTomato reporter knocked into one allele of the endogenous *SFTPC* locus (Hurley et al., 2020), underwent directed differentiation to generate iAT2s in 3D Matrigel cultures as follows. Cells were first differentiated into definitive endoderm using the STEMdiff Definitive Endoderm Kit (Stem Cell Technologies) for 72 hours and subsequently dissociated with GCDR and passaged as small clumps into growth factor reduced Matrigel-coated (Corning) 6-well culture plates (Corning) in ‘‘DS/SB’’ foregut endoderm anteriorization media, consisting of complete serum-free differentiation medium (cSFDM) base as previously described (Jacob et al., 2017), supplemented with 10 µm SB431542 (‘‘SB’’; Tocris) and 2 µm Dorsomorphin (‘‘DS’’; Stemgent), to pattern cells towards anterior foregut endoderm (AFE; day 6 of differentiation). For the first 24 hours after passaging, media was supplemented with 10 µM Y-27632. After anteriorization in DS/SB media for 72 hours, beginning on day 6 of differentiation cells were cultured in ‘‘CBRa’’ lung progenitor-induction medium for 9 additional days. ‘‘CBRa’’ medium consists of cSFDM base supplemented with 3 µM CHIR99021 (Tocris), 10 ng/mL recombinant human BMP4 (rhBMP4, R&D Systems), and 100 nM retinoic acid (RA, Sigma), as described (Jacob et al., 2017). On differentiation day 15, NKX2-1^+^ lung progenitors were isolated based on CD47^hi^/CD26^neg^ gating (Hawkins et al., 2017) using a high-speed cell sorter (MoFlo Legacy or MoFlo Astrios EQ). Purified day 15 lung progenitors were resuspended in undiluted growth factor-reduced 3D Matrigel (Corning) at a concentration of 400 cells/µl and distal/alveolar differentiation was performed in ‘‘CK+DCI’’ medium, consisting of cSFDM base supplemented with 3 µm CHIR99021 (Tocris), 10 ng/mL rhKGF (R&D Systems), and 50 nM dexamethasone (Sigma), 0.1 mM 8-Bromoadenosine 30, 50-cyclic monophosphate sodium salt (Sigma) and 0.1 mM 3-Isobutyl-1-methylxanthine (IBMX; Sigma) (DCI) with a brief period of CHIR99021 withdrawal between days 34-39 to achieve iAT2 maturation. To establish pure cultures of iAT2s, cells were sorted by flow cytometry on day 45 to purify SFTPC^tdTomato+^ cells. iAT2s were maintained as self-renewing monolayered epithelial spheres (“alveolospheres”) through serial passaging every 10-14 days and replating in undiluted growth factor-reduced 3D Matrigel (Corning) droplets at a density of 400 cells/μl in CK+DCI medium, as described (Jacob et al., 2019). iAT2 culture quality and purity was monitored at each passage by flow cytometry, with 95.2 ± 4.2% (mean ± S.D.) of cells expressing SFTPC^tdTomato^ over time, as we have previously detailed (Hurley et al., 2020; Jacob et al., 2017).

Cells at day 6 correspond to the AFG stage and day 261 iAT2s were used for the alveolar stage.

### METHOD DETAILS

#### Generation of *FOXA1*^-/-^, *FOXA2*^-/-^, and *FOXA1/2*^-/-^ H1 hESC lines

To generate homozygous *FOXA1*, *FOXA2*, and FOXA1/2 deletion hESC lines, sgRNAs targeting coding exons within each gene were cloned into Px333-GFP, a modified version of Px333 (Maddalo et al., 2014) which was a gift from Andrea Ventura (Addgene, #64073). The plasmid was transfected into H1 hESCs with XtremeGene 9 (Roche), and 24 hours later 8000 GFP^+^ cells were sorted into a well of six-well plate. Individual colonies that emerged within 5-7 days were subsequently transferred manually into 48-well plates for expansion, genomic DNA extraction, PCR genotyping, and Sanger sequencing. For control clones, the Px333-GFP plasmid was transfected into H1 hESCs, and cells were subjected to the same workflow as H1 hESCs transfected with sgRNAs.

sgRNA oligo used to generate *FOXA1*^-/-^ hESCs: CGCCATGAACAGCATGACTG

sgRNA oligo used to generate *FOXA2*^-/-^ hESCs: CATGAACATGTCGTCGTACG

sgRNA oligos used to generate *FOXA1/2*^-/-^ frameshift hESCs:

*FOXA1*: CGCCATGAACAGCATGACTG
*FOXA2*: CATGAACATGTCGTCGTACG

sgRNA oligos used to generate *FOXA1/2*^-/-^ exon deletion hESCs:

*FOXA1* upstream: GCGACTGGAACAGCTACTAC
*FOXA1* downstream: GCACTGCAATACTCGCCTTA
*FOXA2* upstream: TCCGACTGGAGCAGCTACTA
*FOXA2* downstream: CGGCTACGGTTCCCCCATGC

#### Transduction of CyT49 hESCs with *SCRAM* and sh*PDX1*

To generate shRNA expression vectors, shRNA guide sequences were placed under the control of the human U6 pol III promoter in the pLKO.1-TCR backbone (Moffat et al., 2006), which was a gift from David Root (Addgene, plasmid #10878). Guide sequences were as follows:

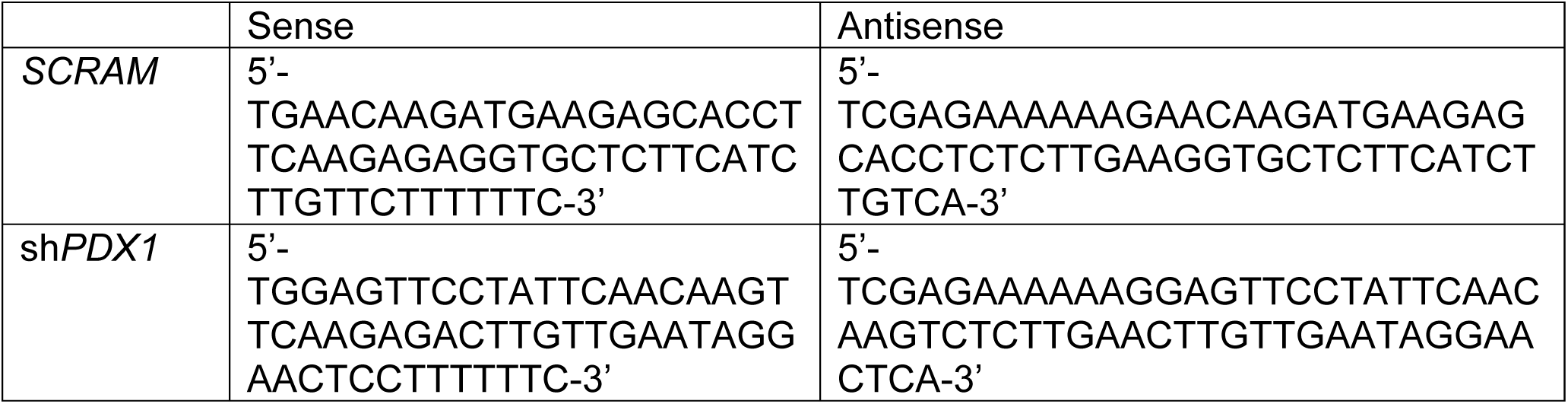

High-titer lentiviral supernatants were generated by co-transfection of the shRNA expression vector and the lentiviral packaging construct into HEK293T cells as described (Xie et al., 2013). Briefly, shRNA expression vectors were co-transfected with the pCMV- R8.74 and pMD2.G expression plasmids (Addgene #22036 and #12259, respectively, gifts from Didier Trono) into HEK293T cells using a 1 mg/ml PEI solution (Polysciences). Lentiviral supernatants were collected at 48 hours and 72 hours after transfection. Lentiviruses were concentrated by ultracentrifugation for 120 min at 19,500 rpm using a Beckman SW28 ultracentrifuge rotor at 4°C.

CyT49 hESCs were plated onto a six-well plate at a density of 1 million cells per well. The following morning, concentrated lentivirus was added at 5 µL/mL media, as well as 8 µg/mL polybrene. After 30 minutes of incubation, the 6 well plate was spun in a centrifuge (Sorvall Legend RT) for 1 hour at 30 C at 950 G. 6 hours later, viral media was replaced with fresh base culture media. After 72 hours, cells were sorted for GFP expression and re-cultured.

#### Immunofluorescence analysis

Cell aggregates derived from hESCs were allowed to settle in microcentrifuge tubes and washed twice with PBS before fixation with 4% paraformaldehyde (PFA) for 30 min at room temperature. Fixed samples were washed twice with PBS and incubated overnight at 4 °C in 30% (w/v) sucrose in PBS. Samples were then loaded into disposable embedding molds (VWR), covered in Tissue-Tek® O.C.T. Sakura® Finetek compound (VWR) and flash frozen on dry ice to prepare frozen blocks. The blocks were sectioned at 10 µm and sections were placed on Superfrost Plus® (Thermo Fisher) microscope slides and washed with PBS for 10 min. Slide-mounted cell sections were permeabilized and blocked with blocking buffer, consisting of 0.15% (v/v) Triton X-100 (Sigma) and 1% (v/v) normal donkey serum (Jackson Immuno Research Laboratories) in PBS, for 1 hour at room temperature. Slides were then incubated overnight at 4 °C with primary antibody solutions. The following day slides were washed five times with PBS and incubated for 1 hour at room temperature with secondary antibody solutions. Cells were washed five times with PBS before coverslips were applied.

All antibodies were diluted in blocking buffer at the ratios indicated below. Primary antibodies used were mouse anti-FOXA1 (1:100 or 1:1000 dilution, Abcam); goat anti-FOXA2 (1:300 dilution, R&D systems); goat anti-SOX17 (1:300 dilution, R&D systems); goat anti-HNF4A (1:1000 dilution, Santa Cruz Biotechnology); rabbit anti-PDX1 (1:500 dilution, Abcam); and mouse anti-NKX6.1 (1:300 dilution, Developmental Studies Hybridoma Bank). Secondary antibodies against mouse, rabbit, and goat were Alexa488- and Cy3-conjugated donkey antibodies (Jackson Immuno Research Laboratories), and were used at dilutions of 1:500 (anti-rabbit Alexa488) or 1:1000 (all other secondary antibodies). Cell nuclei were stained with Hoechst 33342 (1:3000, Invitrogen). Representative images were obtained with a Zeiss Axio-Observer-Z1 microscope equipped with a Zeiss ApoTome and AxioCam digital camera. Figures were prepared in Adobe Creative Suite 5.

#### Flow cytometry analysis

Cell aggregates derived from hESCs were allowed to settle in microcentrifuge tubes and washed with PBS. Cell aggregates were incubated with Accutase® at room temperature until a single-cell suspension was obtained. Cells were washed with 1 mL ice-cold flow buffer comprised of 0.2% BSA in PBS and centrifuged at 200 g for 5 min. BD Cytofix/Cytoperm™ Plus Fixation/Permeabilization Solution Kit was used to fix and stain cells for flow cytometry according to the manufacturer’s instructions. Briefly, cell pellets were re-suspended in ice-cold BD Fixation/Permeabilization solution (300 µL per microcentrifuge tube). Cells were incubated for 20 min at 4 °C. Cells were washed twice with 1 mL ice-cold 1X BD Perm/Wash™ Buffer and centrifuged at 10 °C and 200 x g for 5 min. Cells were re-suspended in 50 µL ice-cold 1X BD Perm/Wash™ Buffer containing diluted antibodies, for each staining performed. Cells were incubated at 4 °C in the dark for 1-3 hours. Cells were washed with 1.25 mL ice-cold 1X BD Wash Buffer and centrifuged at 200 g for 5 min. Cell pellets were re-suspended in 300 µL ice-cold flow buffer and analysed in a FACSCanto™ II (BD Biosciences). Antibodies used were PE-conjugated anti-SOX17 antibody (1:20 dilution, BD Biosciences); mouse anti-HNF1B antibody (1:100 dilution, Santa Cruz Biotechnology); PE-conjugated anti-mouse IgG (1:50 dilution, BD Biosciences); PE-conjugated anti-PDX1 (1:10 dilution, BD Biosciences); and AlexaFluor® 647-conjugated anti-NKX6.1 (1:5 dilution, BD Biosciences). Data were processed using FlowJo software v10.

#### Chromatin Immunoprecipitation Sequencing (ChIP-seq)

ChIP-seq was performed using the ChIP-IT High-Sensitivity kit (Active Motif) according to the manufacturer’s instructions. Briefly, for each cell stage and condition analyzed, 5-10 x 10^6^ cells were harvested and fixed for 15 min in an 11.1% formaldehyde solution. Cells were lysed and homogenized using a Dounce homogenizer and the lysate was sonicated in a Bioruptor® Plus (Diagenode), on high for 3 x 5 min (30 sec on, 30 sec off). Between 10 and 30 µg of the resulting sheared chromatin was used for each immunoprecipitation. Equal quantities of sheared chromatin from each sample were used for immunoprecipitations carried out at the same time. 4 µg of antibody were used for each ChIP-seq assay. Chromatin was incubated with primary antibodies overnight at 4 °C on a rotator followed by incubation with Protein G agarose beads for 3 hours at 4 °C on a rotator. Antibodies used were rabbit anti-H3K27ac (Active Motif 39133); rabbit anti-H3K4me1 (Abcam ab8895); goat anti-FOXA1 (Abcam Ab5089); goat-anti-FOXA2 (Santa Cruz SC-6554); and mouse anti-HNF4A (Novus PP-H1415). Reversal of crosslinks and DNA purification were performed according to the ChIP-IT High-Sensitivity instructions, with the modification of incubation at 65 °C for 2-3 hours, rather than at 80 °C for 2 hours. Sequencing libraries were constructed using KAPA DNA Library Preparation Kits for Illumina® (Kapa Biosystems) and library sequencing was performed on either a HiSeq 4000 System (Illumina®) or NovaSeq 6000 System (Illumina®) with single-end reads of either 50 or 75 base pairs (bp). Sequencing was performed by the UCSD Institute for Genomic Medicine (IGM) core research facility. For ChIP-seq experiments at the DE, AFG, and ALV stages in iAEC2 cells, two technical replicates from a single differentiation were generated. For all other ChIP-seq experiments, replicates from two independent hESC differentiations were generated.

#### ChIP-seq data analysis

ChIP-seq reads were mapped to the human genome consensus build (hg19/GRCh37) and visualized using the UCSC Genome Browser (Kent et al., 2002). Burrows-Wheeler Aligner (BWA) (Li and Durbin, 2009) version 0.7.13 was used to map data to the genome. Unmapped and low-quality (q<15) reads were discarded. SAMtools (Li et al., 2009) version 1.5 was used to remove duplicate sequences and HOMER (Heinz et al., 2010) version 4.10.4 was used to call peaks using the findPeaks command with default parameters. The command “-style factor” was used for TFs and the command “-style histone” was used for histone modifications. Stage- and condition-matched input DNA controls were used as background when calling peaks. The BEDtools (Quinlan and Hall, 2010) version 2.26.0 suite of programs was used to perform genomic algebra operations. Tag directories were created for each replicate using HOMER. Directories from each replicate were then combined, and peaks were called from the combined replicates using HOMER. These peaks were then intersected with pancreatic enhancers, hepatic enhancers, or alveolar enhancers, respectively. Pearson correlations for the intersecting peaks were calculated between each pair of replicates using the command multiBamSummary from the deepTools2 package (Ramirez et al., 2016) version 3.1.3 and are as follows:

|  | H3K27ac | H3K4me1 | FOXA1 | FOXA2 | PDX1 | HNF6 | SOX9 | HNF4A |
| --- | --- | --- | --- | --- | --- | --- | --- | --- |
| CyT49 ES | 0.94 | N/A | N/A | N/A | N/A | N/A | N/A | N/A |
| CyT49 DE | 0.91 | N/A | N/A | N/A | N/A | N/A | N/A | N/A |
| CyT49 GT | 0.88 | 0.98 | 0.90 | 0.90 | N/A | N/A | N/A | N/A |
| CyT49 PP1 | 0.83 | N/A | N/A | N/A | N/A | N/A | N/A | N/A |
| CyT49 PP2 | 0.77 | 0.99 | 0.89 | 0.84 | 0.64 | 0.93 | 0.86 | N/A |
| CyT49 HP | 0.94 | N/A | 0.84 | 0.70 | N/A | N/A | N/A | 0.88 |
| CyT49 PP2<br>*SCRAM* | 0.71 | N/A | 0.90 | 0.68 | N/A | N/A | N/A | N/A |
| CyT49 PP2<br>*shPDX1* | 0.86 | N/A | 0.86 | 0.62 | N/A | N/A | N/A | N/A |
| H1 GT | 0.98 | 0.86 | N/A | N/A | N/A | N/A | N/A | N/A |
| H1 PP2 | 0.97 | 0.88 | N/A | N/A | N/A | N/A | N/A | N/A |
| H1<br>*FOXA1/2*<br>exon<br>deletion<br>GT | 0.98 | 0.87 | N/A | N/A | N/A | N/A | N/A | N/A |
| H1<br>*FOXA1/2*<br>exon<br>deletion<br>PP2 | 0.99 | 0.88 | N/A | N/A | N/A | N/A | N/A | N/A |
| iAEC2 DE | 1.00 | N/A | N/A | N/A | N/A | N/A | N/A | N/A |
| iAEC2<br>AFG | 1.00 | N/A | 0.99 | N/A | N/A | N/A | N/A | N/A |
| iAEC2<br>ALV | 1.00 | N/A | 0.95 | N/A | N/A | N/A | N/A | N/A |

#### RNA isolation and sequencing (RNA-seq) and qRT-PCR

RNA was isolated from cell samples using the RNeasy® Micro Kit (Qiagen) according to the manufacturer instructions. For each cell stage and condition analyzed between 0.1 and 1 x 10^6^ cells were collected for RNA extraction. For qRT-PCR, cDNA synthesis was first performed using the iScript™ cDNA Synthesis Kit (Bio-Rad) and 500 ng of isolated RNA per reaction. qRT-PCR reactions were performed in triplicate with 10 ng of template cDNA per reaction using a CFX96™ Real-Time PCR Detection System and the iQ™ SYBR® Green Supermix (Bio-Rad). PCR of the TATA binding protein (TBP) coding sequence was used as an internal control and relative expression was quantified via double delta CT analysis. For RNA-seq, stranded, single-end sequencing libraries were constructed from isolated RNA using the TruSeq® Stranded mRNA Library Prep Kit (Illumina®) and library sequencing was performed on either a HiSeq 4000 System (Illumina®) or NovaSeq 6000 System (Illumina®) with single-end reads of either 50 or 75 base pairs (bp). Sequencing was performed by the UCSD IGM core research facility. A complete list of RT-qPCR primer sequences can be found below.

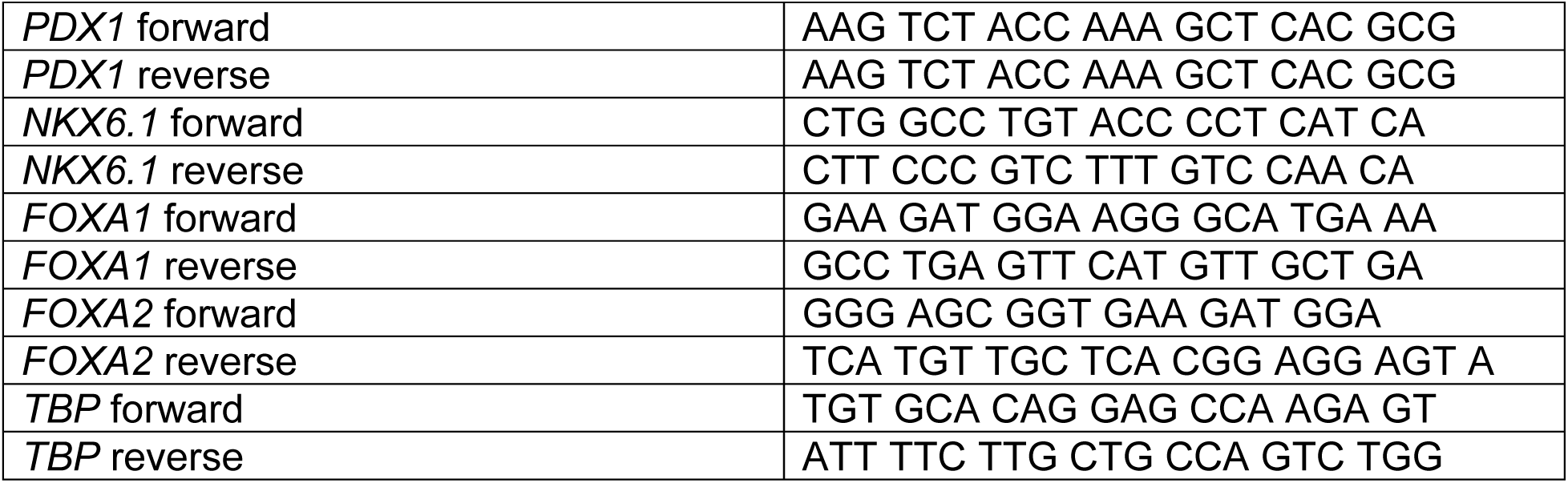

#### RNA-seq data analysis

Reads were mapped to the human genome consensus build (hg19/GRCh37) using the Spliced Transcripts Alignment to a Reference (STAR) aligner version 2.4 (Dobin et al., 2013). Normalized gene expression (fragments per kilobase per million mapped reads; FPKM) for each sequence file was determined using Cufflinks (Trapnell et al., 2010) version 2.2.1 with the parameters: --library-type fr-firststrand --max-bundle-frags 10000000. Differential gene expression was determined using DESeq2 (Love et al., 2014). Adjusted *P-*values < 0.05 and fold change ≥ 2 were considered significant. For RNA-seq corresponding to cells at the HP stage, one replicate was generated. For all other RNA-seq experiments, replicates from two independent hESC differentiations were generated. Pearson correlations between bam files corresponding to each pair of replicates were calculated, and are as follow:

|  | Non-transfected<br>control H1<br>hESCs | Non-edited<br>H1 hESCs | *FOXA1/2* frameshift<br>mutation H1 hESCs | *FOXA1/2*<br>exon deletion<br>H1 hESCs |
| --- | --- | --- | --- | --- |
| DE | 0.89 | 0.92 | 0.92 | 0.93 |
| GT | 0.96 | 0.91 | 0.91 | 0.92 |
| PP1 | 0.98 | 0.98 | 0.93 | 0.94 |
| PP2 | 0.99 | 0.96 | 0.83 | 0.90 |

#### Assay for Transposase Accessible Chromatin Sequencing (ATAC-seq)

ATAC-seq (Buenrostro et al., 2013) was performed on approximately 50,000 nuclei. The samples were permeabilized in cold permabilization buffer (0.2% IGEPAL-CA630 (I8896, Sigma), 1 mM DTT (D9779, Sigma), Protease inhibitor (05056489001, Roche), 5% BSA (A7906, Sigma) in PBS (10010-23, Thermo Fisher Scientific) for 10 minutes on the rotator in the cold room and centrifuged for 5 min at 500 xg at 4°C. The pellet was resuspended in cold tagmentation buffer (33 mM Tris-acetate (pH = 7.8) (BP-152, Thermo Fisher Scientific), 66 mM K-acetate (P5708, Sigma), 11 mM Mg-acetate (M2545, Sigma), 16% DMF (DX1730, EMD Millipore) in Molecular biology water (46000-CM, Corning)) and incubated with tagmentation enzyme (FC-121-1030; Illumina) at 37 °C for 30 min with shaking at 500 rpm. The tagmented DNA was purified using MinElute PCR purification kit (28004, QIAGEN). Libraries were amplified using NEBNext High-Fidelity 2X PCR Master Mix (M0541, NEB) with primer extension at 72°C for 5 minutes, denaturation at 98°C for 30 s, followed by 8 cycles of denaturation at 98°C for 10 s, annealing at 63°C for 30 s and extension at 72°C for 60 s. After the purification of amplified libraries using MinElute PCR purification kit (28004, QIAGEN), double size selection was performed using SPRIselect bead (B23317, Beckman Coulter) with 0.55X beads and 1.5X to sample volume. Finally, libraries were sequenced on HiSeq4000 (Paired-end 50 cycles, Illumina).

#### ATAC-seq data analysis

ATAC-seq reads were mapped to the human genome (hg19/GRCh37) using Burrows-Wheeler Aligner (Li and Durbin, 2009) (BWA) version 0.7.13, and visualized using the UCSC Genome Browser (Kent et al., 2002). SAMtools (Li et al., 2009) was used to remove unmapped, low-quality (q<15), and duplicate reads. MACS2 (Zhang et al., 2008) version 2.1.4 was used to call peaks, with parameters “shift set to 100 bps, smoothing window of 200 bps” and with “nolambda” and “nomodel” flags on. MACS2 was also used to call ATAC-Seq summits, using the same parameters combined with the “call-summits” flag.

For all ATAC-seq experiments, replicates from two independent hESC differentiations were generated. Bam files for each pair of replicates were merged for downstream analysis using SAMtools, and Pearson correlations between bam files for each individual replicate were calculated over a set of peaks called from the merged bam file. Correlations were performed using the command multiBamSummary from the deepTools2 package (Ramirez et al., 2016) with the “--removeOutliers” flag and are as follows:

|  | Non-transfected<br>control H1<br>hESCs | *FOXA1/2* exon<br>deletion H1<br>hESCs | *SCRAM*<br>transduced<br>CyT49 hESCs | sh*PDX1*<br>transduced<br>CyT49 hESCs |
| --- | --- | --- | --- | --- |
| GT | 0.99 | 0.98 | N/A | N/A |
| PP2 | 0.97 | 0.99 | 0.99 | 1.00 |

#### Hi-C data analysis

Hi-C data were processed as previously described with some modifications (Dixon et al., 2015). Read pairs were aligned to the hg19 reference genome separately using BWA-MEM with default parameters (Li and Durbin, 2009). Specifically, chimeric reads were processed to keep only the 5’ position and reads with low mapping quality (<10) were filtered out. Read pairs were then paired using custom scripts. Picard tools were then used to remove PCR duplicates. Bam files with alignments were further processed into text format as required by Juicebox tools (Durand et al., 2016). Juicebox tools were then applied to generate Hi-C files containing normalized contact matrices. All downstream analysis was based on 10 Kb resolution KR normalized matrices.

Chromatin loops were identified by comparing each pixel with its local background, as described previously (Rao et al., 2014) with some modifications. Specifically, only the donut region around the pixel was compared to model the expected count. Briefly, the KR-normalized contact matrices at 10 Kb resolution were used as input for loop calling. For each pixel, distance-corrected contact frequencies were calculated for each surrounding bin and the average of all surrounding bins. The expected counts were then transformed to raw counts by multiplying the counts with the raw-to-KR normalization factor. The probability of observing raw expected counts was calculated using Poisson distribution. All pixels with *P-*value < 0.01 and distance less than 10 Kb were selected as candidate pixels. Candidate pixels were then filtered to remove pixels without any neighboring candidate pixels since they were likely false positives. Finally, pixels within 20 Kb of each other were collapsed and only the most significant pixel was selected. The collapsed pixels with *P*-value < 1 x 10^-5^ were used as the final list of chromatin loops.

#### Gene Ontology analysis

Gene ontology analysis for enhancer groups was performed using GREAT (McLean et al., 2010) version 4.0.4 with the default parameters. Gene ontology for differentially expressed genes was performed using Metascape (Zhou et al., 2019) using default parameters.

#### Identification of super-enhancers

Super-enhancers were defined using the Rank Order Super-Enhancers (ROSE) package (Loven et al., 2013; Whyte et al., 2013) using default parameters. Specifically, pancreatic enhancers were ranked based on PP2 H3K27ac signal and super-enhancers were defined as enhancers ranked beyond the inflection point.

#### Principle component analysis

For RNA-seq data, transcriptomes were first filtered for genes expressed (FPKM ≥ 1) in at least one condition, then log10 transformed. For distal H3K27ac signals, H3K27ac peaks were filtered for distal enhancers (≥ 2.5 kb from any annotated TSS). Based on filtered values, PCA plots were generated using the PRComp package in R.

#### Quantification of changes in H3K27ac signal

HOMER (Heinz et al., 2010) was used to annotate raw H3K27ac ChIP-seq reads over distal enhancers at developmental stages both before and after lineage induction. HOMER was then used to invoke the R package DESeq2 (Love et al., 2014) version 3.10 for differential analysis, using default parameters.

#### Quantification of changes in TF ChIP-seq and ATAC-seq signal

HOMER (Heinz et al., 2010) was used to annotate raw FOXA1 and FOXA2 ChIP-seq reads, as well as ATAC-seq reads over PDX1-bound class I and class II enhancers in cells transfected with *SCRAM* and sh*PDX1* lentivirus. HOMER was then used to invoke the R package DESeq2 (Love et al., 2014) for differential analysis, using the flag “norm2total.”

#### Assignment of enhancer target genes

RNA-seq data were filtered for expressed genes (FPKM ≥ 1) at the PP2 stage, and BEDTools (Quinlan and Hall, 2010) “closest” command was used to assign each enhancer to the nearest annotated TSS.

#### Motif enrichment analysis

HOMER (Heinz et al., 2010) was used for comparative motif enrichment analyses, using the command findMotifsGenome.pl. *de novo* motifs were assigned to TFs based on suggestions generated by HOMER.

#### Identification of FOXA motifs and generation of log-odds scores

FOXA1 and FOXA2 PWMs were downloaded from the JASPAR database (Fornes et al., 2020), and occurrences with associated log-odds scores were quantified using the FIMO feature within the MEMEsuit package (Grant et al., 2011) version 5.1.1.

#### Calculation of positional motif enrichment

Identified ATAC-seq summits on class I and class II enhancers were flanked by 500 bp in each direction, and the CENTRIMO feature within the MEMEsuit package (Bailey and Machanick, 2012) version 5.1.1 was used to determine enrichment at summits for selected PWMs associated with FOXA1 and FOXA2, as well as to graph the positional probability of motif occurrence with respect to ATAC-seq summits.

#### ATAC-seq footprinting analysis

ATAC-seq footprinting was performed as previously described (Aylward et al., 2018). In brief, diploid genomes for CyT49 were created using vcf2diploid (version 0.2.6a) (Rozowsky et al., 2011) and genotypes called from whole genome sequencing and scanned for a compiled database of TF sequence motifs from JASPAR (Mathelier et al., 2016) and ENCODE (Consortium, 2012) with FIMO (Grant et al., 2011) using default parameters for *P*-value threshold and a 40.9% GC content based on the hg19 human reference genome. Footprints within ATAC-seq peaks were discovered with CENTIPEDE (version 1.2) (Pique-Regi et al., 2011) using cut-site matrices containing Tn5 integration counts within a ±100 bp window around each motif occurrence. Footprints were defined as those with a posterior probability ≥ 0.99.

#### Permutation-based significance

A random sampling approach (10,000 iterations) was used to obtain null distributions for enrichment analyses, in order to obtain *P*-values. Null distributions for enrichments were obtained by randomly shuffling enhancer regions using BEDTools (Quinlan and Hall, 2010) and overlapping with FOXA1/2 binding sites. *P*-values < 0.05 were considered significant.

#### Data sources

The following datasets used in this study were obtained from the GEO and ArrayExpress repositories:

RNA-seq: Pancreatic differentiation of CyT49 hESC line (E-MTAB-1086)

ChIP-seq: H3K27ac in CyT49 hESC, DE, GT, PP1, PP2 (GSE54471 and GSE149148); H3K27ac in CyT49 PP2 SCRAM and PP2 sh*PDX1* (GSE54471); H3K4me1 in CyT49 GT and PP2 (GSE54471 and GSE149148); PDX1 in CyT49 PP2 (GSE54471 and GSE149148); HNF6 in CyT49 PP2 (GSE149148); SOX9 in CyT49 PP2 (GSE149148); FOXA1 in CyT49 PP2 (GSE149148); FOXA2 in CyT49 PP2 (GSE149148).

ATAC-seq: CyT49 GT and PP2 (GSE149148)

Hi-C datasets were generated as a component of the 4D Nucleome Project (Dekker et al., 2017). Datasets corresponding to the PP2 stages of differentiation can be found under accession number 4DNES0LVRKBM.

### QUANTIFICATION AND STATISTICAL ANALYSES

Statistical analyses were performed using GraphPad Prism (v8.1.2), and R (v3.6.1). Statistical parameters such as the value of n, mean, standard deviation (SD), standard error of the mean (SEM), significance level (n.s., not significant; *p < 0.05; **p < 0.01; and ***p < 0.001), and the statistical tests used are reported in the figures and figure legends. The ‘‘n’’ refers to the number of independent hESC differentiation experiments analyzed (biological replicates). All bar graphs and line graphs are displayed as mean ± S.E.M, and all box plots are centered on median, with box encompassing 25th-75th percentile and whiskers extending up to 1.5 interquartile range. Statistically significant gene expression changes were determined with DESeq2 (Love et al., 2014).

### DATA AVAILABILITY

All mRNA-seq, ChIP-seq, and ATAC-seq datasets generated for this study have been deposited at GEO under the accession number GSE148368.

## ACKNOWLEDGEMENTS

We thank Ileana Matta for assistance with ATAC-seq assays and library preparations, and members of the Sander laboratory, Dr. Christopher W. Benner, Dr. Emma Farley, and Dr. Xin Sun for helpful discussions and critical reading of the manuscript. We acknowledge support of the UCSD Human Embryonic Stem Cell Core for cell sorting and K. Jepsen and the UCSD IGM Genomic Center (supported by P30 DK063491) for library preparation and sequencing. This work was supported by grant T32 GM008666 (R.J.G.), R01 DK068471 and R01 DK078803 (M.S.), U54 DK107977 (B.R.), the I.M. Rosenzweig Junior Investigator Award from the Pulmonary Fibrosis Foundation (K.D.A.), R01 HL095993, R01HL128172, U01TR001810, and N01 75N92020C00005 (D.N.K).

## AUTHOR CONTRIBUTIONS

A.W. and M.S. conceived the project. R.J.G, A.W., and M.S. designed experiments. A.W., R.J.G., D.K.L, N.K.V, D.A.R, K.D.A, J.W., and S.K. performed experiments. A.W., R.J.G, N.K.V., Y.Q., and J.C. analyzed sequencing data. R.J.G, A.W, and M.S. interpreted data. R.J.G. and M.S. wrote the manuscript. B.R., K.J.G., D.N.K., and M.S. supervised all research.

---

**Figure S1, related to Figure 1.**
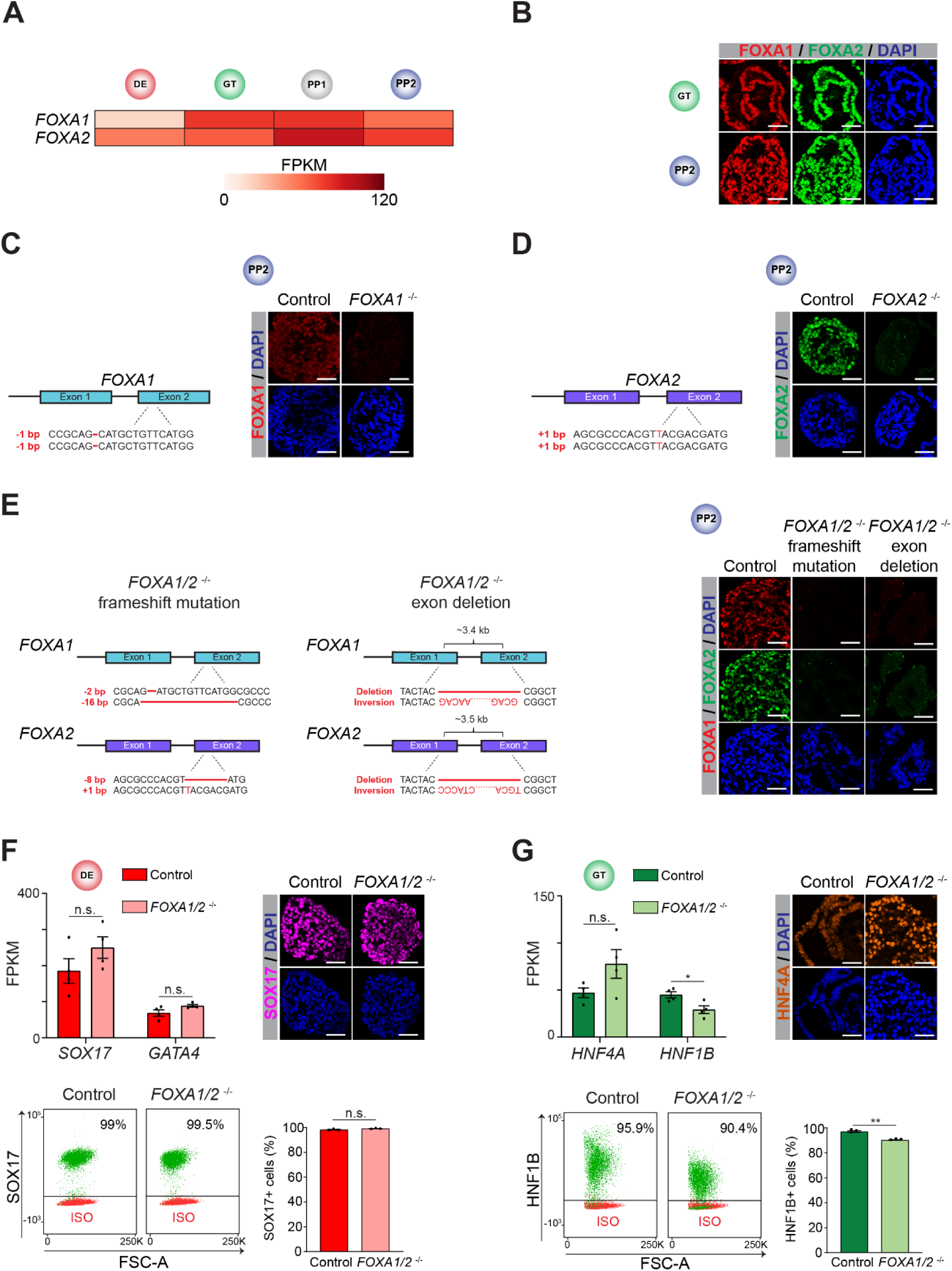
Definitive endoderm and gut tube specification does not require FOXA1 and FOXA2. (**A**) Heatmap showing mRNA expression levels of *FOXA1* and *FOXA2* determined by RNA-seq during pancreatic differentiation of hESCs. FPKM, Fragments per kilobase per million fragments mapped. (**B**) Representative immunofluorescent staining of FOXA1 and FOXA2 in GT and PP2. (**C**) Schematic of frameshift mutation in *FOXA1*^-/-^ hESCs (left) and representative immunofluorescent staining (right) of FOXA1 in PP2. (**D**) Schematic of frameshift mutation in *FOXA2*^-/-^ hESCs (left) and representative immunofluorescent staining (right) of FOXA2 in PP2. (**E**) Schematic of frameshift mutations and exon deletions in *FOXA1/2*^-/-^ hESCs (left) and representative immunofluorescent staining (right) of FOXA1 and FOXA2 in PP2. (**F** and **G**) mRNA expression levels determined by RNA-seq (left), representative immunofluorescent staining (right), and flow cytometry analysis and quantification (bottom) in control and *FOXA1/2*^-/-^ DE (**F**) and GT (**G**) cells (n = 4 and n = 3 independent differentiations for RNA-seq and flow cytometry, respectively; qPCR: *P* adj. = 0.116 and 0.104 for *SOX17* and *GATA4*, respectively, in **F** and 0.014 and 0.061 for *HNF1B* and *HNF4A*, respectively, in **G**; DESeq2; n.s., not significant; flow cytometry: *P* = 0.116 in control compared to *FOXA1/2*^-/-^ DE cells in **F** and 4.5 x 10^-3^ in control compared to *FOXA1/2*^-/-^ GT cells in **G**; student’s t-test, 2 sided). FSC-A, forward scatter area. For all immunofluorescence, representative images are shown from n ≥ 2 independent differentiations. Scale bars, 50 µm. For all flow cytometry analyses, representative plots are shown from n = 3 independent differentiations, with isotype control (ISO) for each antibody shown in red and target protein staining in green.

**Figure S2, related to Figure 2.**
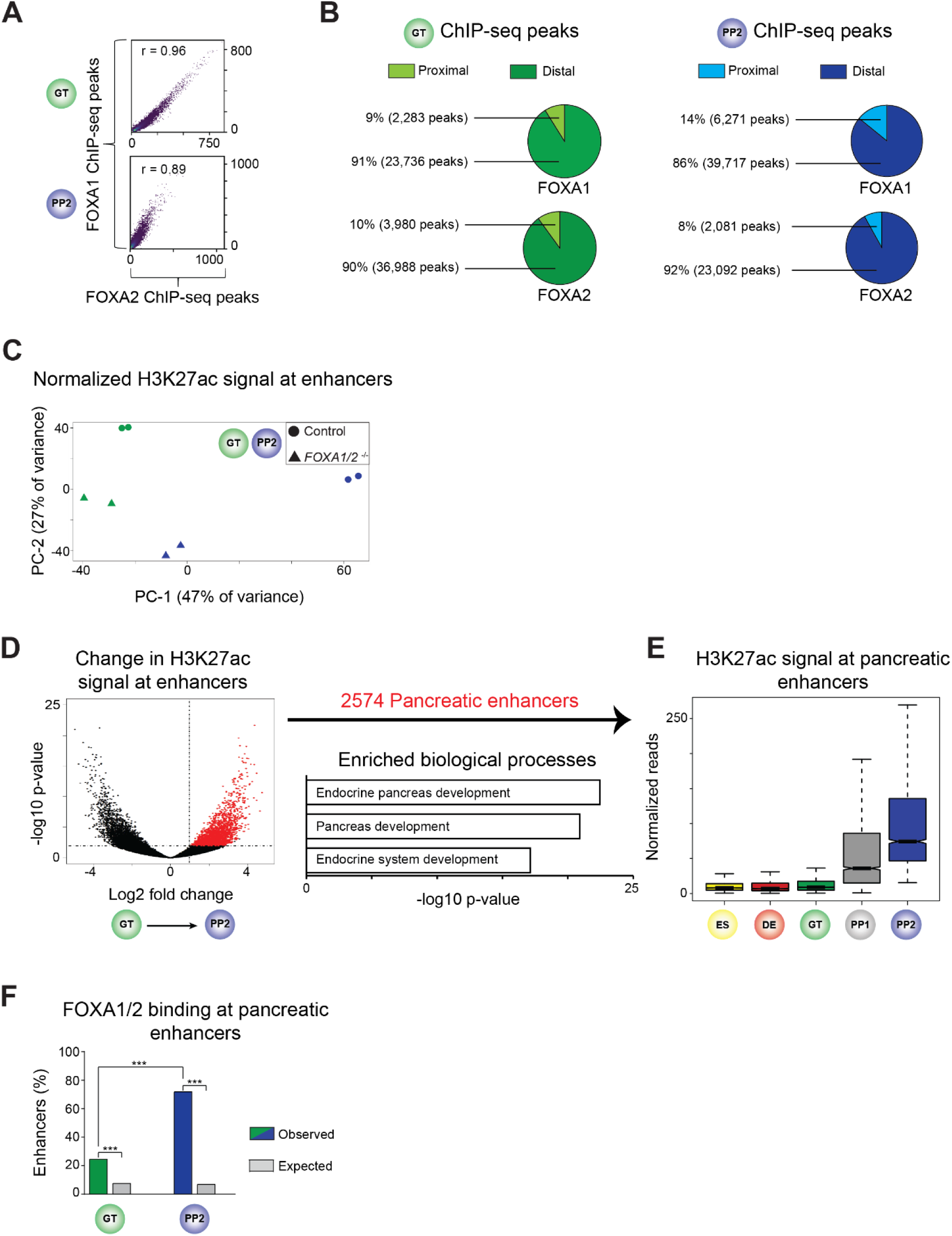
FOXA1 and FOXA2 both bind to pancreas-specific enhancers. (**A**) Pearson correlation between FOXA1 and FOXA2 ChIP-seq signal at FOXA1 and FOXA2 peaks in GT and PP2. (**B**) Percentage of FOXA1 and FOXA2 peaks located proximal (≤ 2.5 kb) or distal (> 2.5 kb) to nearest annotated TSS. (**C**) Principle component analysis showing variance in distal (> 2.5 kb from TSS) H3K27ac signal between control and *FOXA1/2*^-/-^ cells in GT and PP2. Each plotted point represents one biological replicate. (**D**) Volcano plot showing identification of pancreatic enhancers based on increase in H3K27ac signal from GT to PP2 (≥ 2-fold increase, *P* adj. < 0.05 at sites > 2.5 kb from TSS). Enriched gene ontology terms of genes linked to pancreatic enhancers using GREAT. (**E**) Box plots of H3K27ac ChIP-seq counts at pancreatic enhancers. (**F**) Enrichment of pancreatic enhancers for FOXA1 or FOXA2 peaks compared to random genomic regions at GT and PP2 (*P* < .0001 and *P* < .0001, respectively; permutation test). Pancreatic enhancers are enriched for FOXA1/2 peaks at PP2 compared to GT (*P* < 2.2 x 10^-16^, Fisher’s exact test, 2-sided). All ChIP-seq experiments, n = 2 replicates from independent differentiations.

**Figure S3, related to Figure 3.**
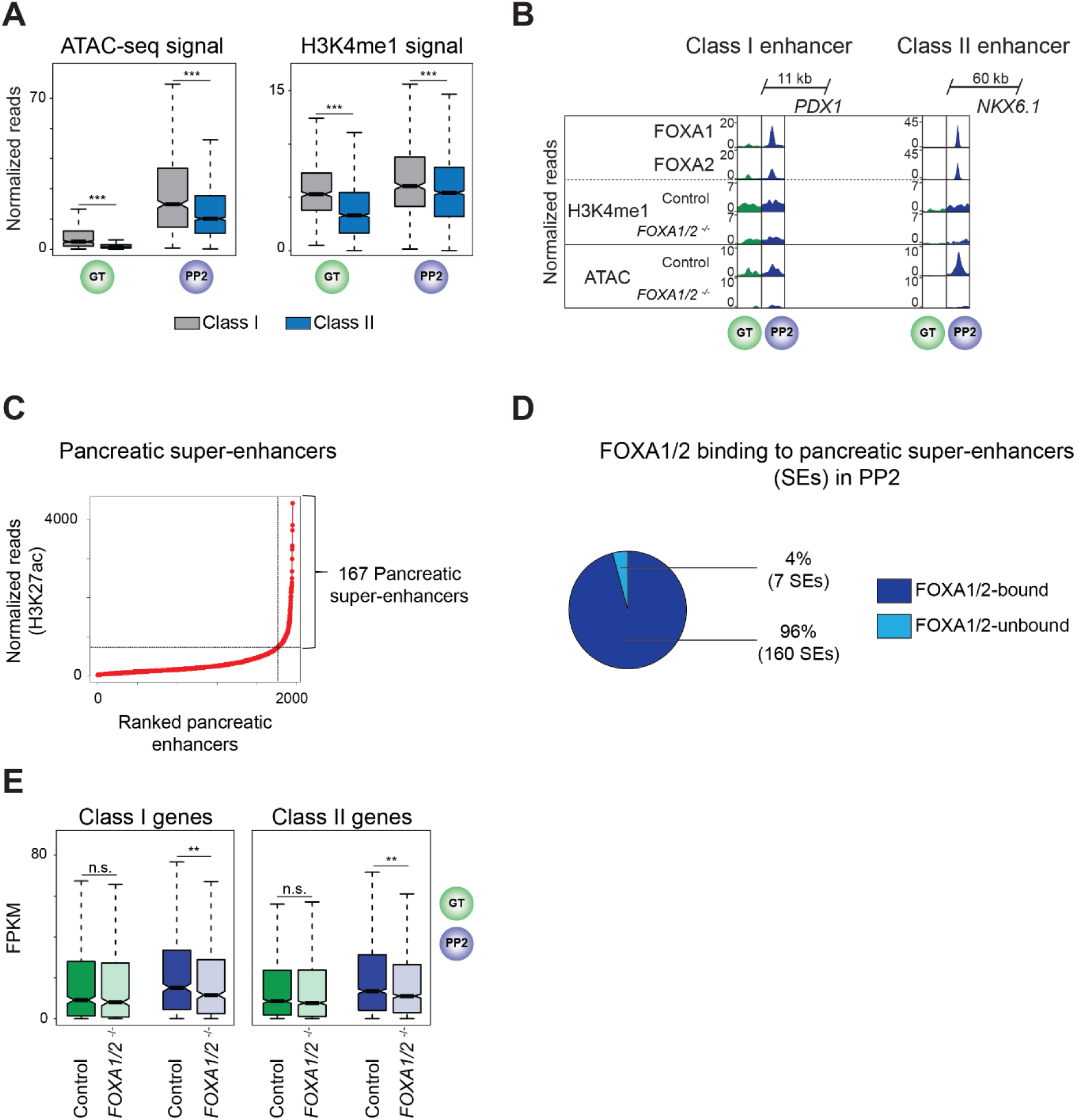
Characterization of class I and class II pancreatic enhancers. (**A**) Box plots of ATAC-seq and H3K4me1 ChIP-seq counts at class I and class II enhancers in GT and PP2 (*P* = < 2.2 x 10^-16^, 1.11 x 10^-14^, < 2.2 x 10^-16^, and 1.07 x 10^-7^ for comparisons of ATAC-seq signal in GT and PP2 and comparisons of H3K4me1 ChIP-seq signal in GT and PP2, respectively; Wilcoxon rank sum test, 2- sided). (**B**) Genome browser snapshots showing FOXA1, FOXA2, H3K4me1 ChIP-seq, and ATAC-seq signal at class I enhancer near *PDX1* and class II enhancer near *NKX6.1* in GT and PP2 cells. Approximate distance between enhancer and gene body is indicated. (**C**) Identification of pancreatic super-enhancers by ranking 2574 pancreatic enhancers based on H3K27ac ChIP-seq signal in PP2. (**D**) Percentage of pancreatic super-enhancers containing FOXA1 and/or FOXA2 ChIP-seq peaks in PP2. (**E**) Box plots of mRNA levels (FPKM, fragments per kilobase per million fragments mapped) in control and *FOXA1/2*^-/-^ GT and PP2 cells for genes linked to pancreatic class I and class II enhancers (n = 4 independent differentiations; *P* = 0.182, 4.82 x 10^-3^, 0.067, and 1.21 x 10^-3^ for control versus *FOXA1/2*^-/-^ of class I genes in GT, class I genes in PP2, class II genes in GT, and class II genes in PP2, respectively; Wilcoxon rank sum test, 2-sided). All ChIP-seq and ATAC-seq experiments, n = 2 replicates from independent differentiations.

**Figure S4, related to Figure 4.**
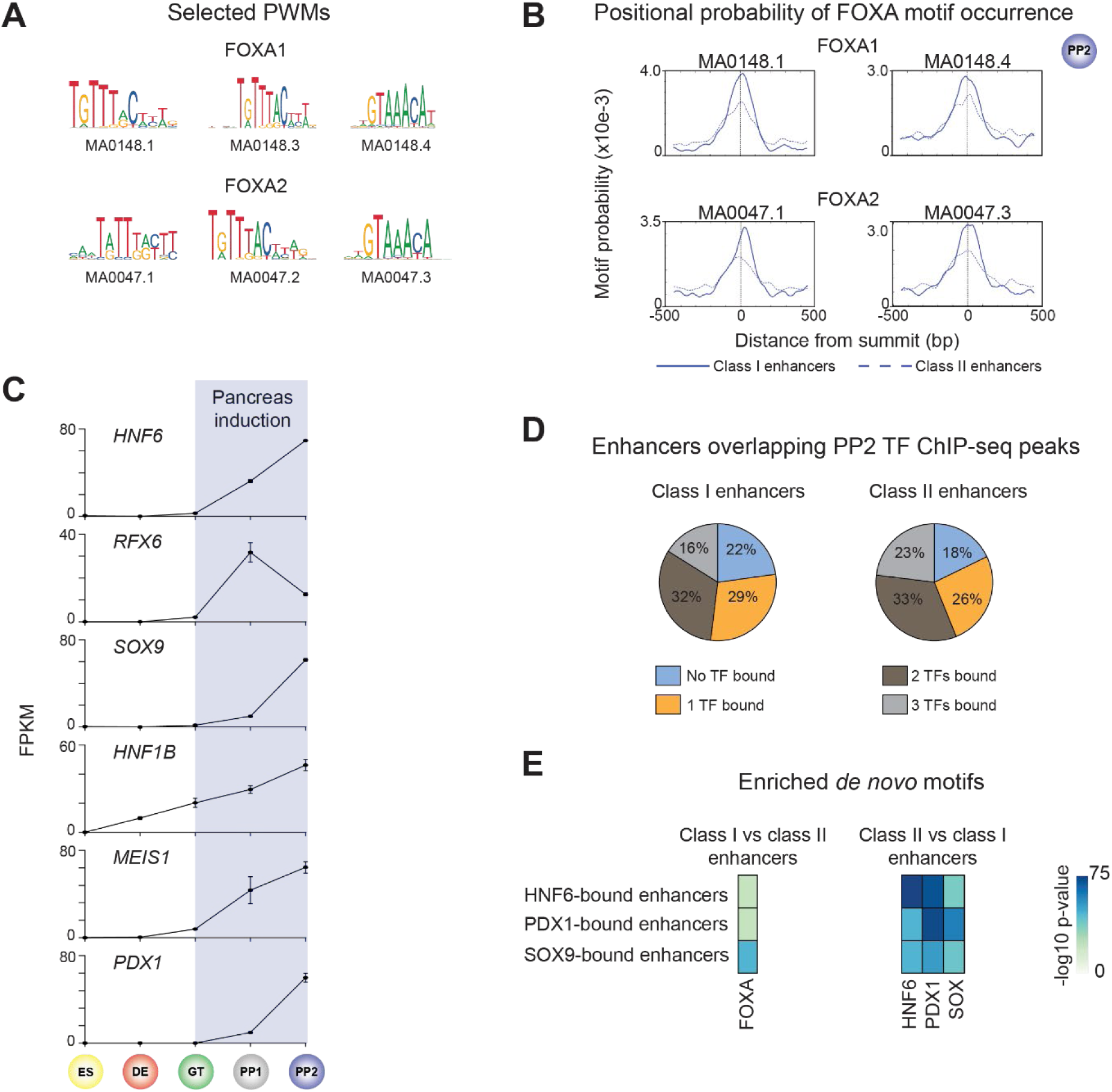
Class I and class II pancreatic enhancers exhibit distinct enhancer architecture. (**A**) Selected FOXA1 and FOXA2 motifs and associated position weight matrices (PWMs) obtained from JASPAR. (**B**) Probability (occurrence per base pair) of FOXA1 (MA0148.1 and MA0148.4) and FOXA2 (MA0047.1 and MA0047.3) motifs relative to ATAC-seq peak summits at class I (solid line) and class II (dashed line) enhancers. ATAC-seq peak summits at class I enhancers are enriched for occurrences compared to summits at class II enhancers (*P* = 8.4 x 10^-15^, 1.6 x 10^-3^, 1.3 x 10^-3^, and 2.1 x 10^-5^ for MA0148.1, MA0148.4, MA0047.1, and MA0047.3, respectively; Fisher’s exact test, 1-sided). (**C**) mRNA expression levels of pancreatic transcription factors (TF) determined by RNA-seq. Data are shown as mean fragments per kilobase per million fragments mapped (FPKM) ± S.E.M. in ES, DE, GT, PP1, and PP2 (n = 3 independent differentiations). (**D**) Percentage of class I and class II enhancers overlapping HNF6, PDX1, and SOX9 ChIP-seq peaks (within 100 bp from peak) in PP2. (**E**) Heatmap showing enriched *de novo* TF binding motifs at HNF6-, PDX1-, and SOX9-bound class I against a background of HNF6-, PDX1-, and SOX9-bound class II enhancers and vice versa. Fisher’s exact test, 1-sided, corrected for multiple comparisons.

**Figure S5, related to Figure 5.**
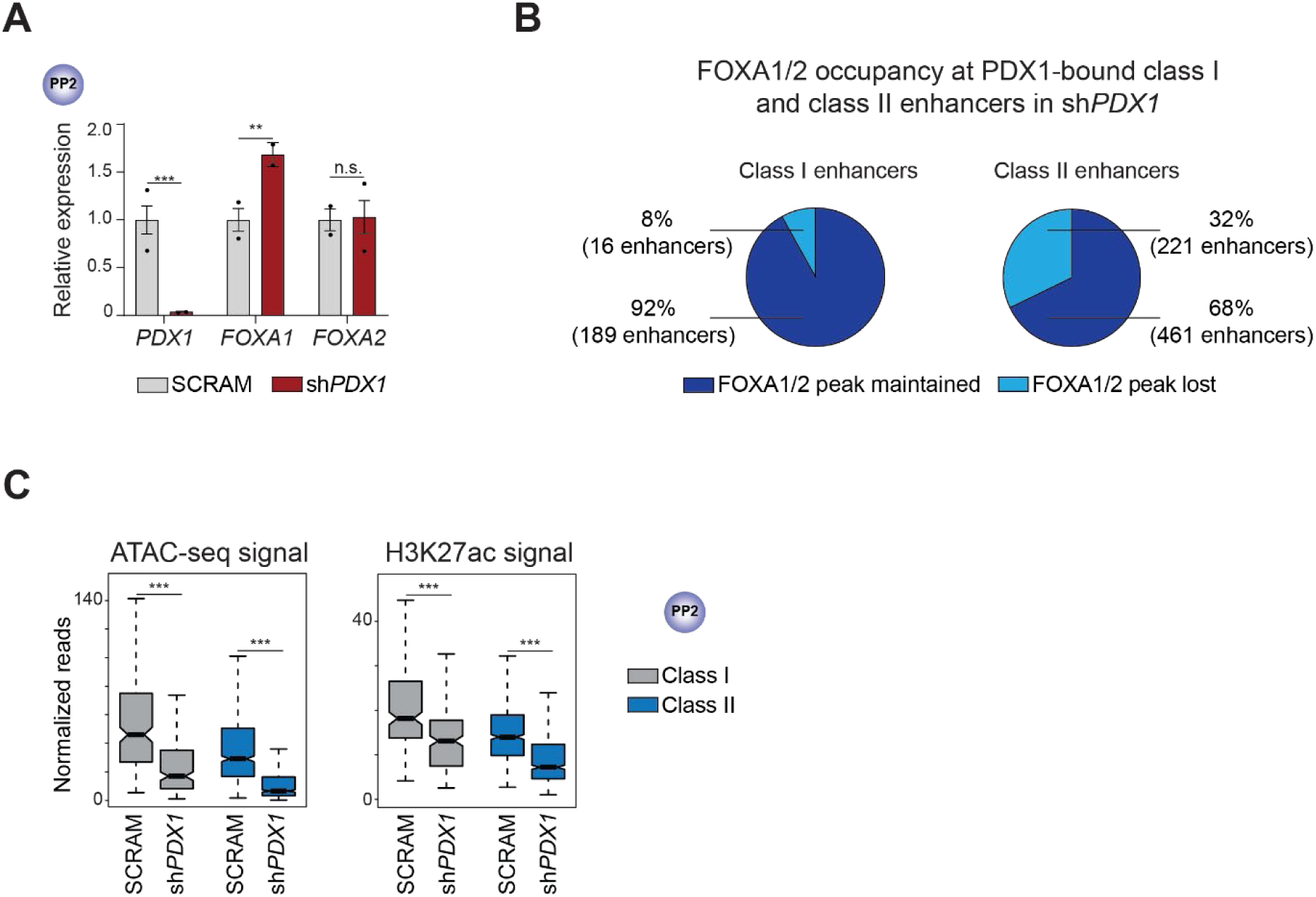
Characterization of class I and class II enhancers in PDX1-deficient pancreatic progenitors. (**A**) qPCR analysis of *PDX1*, *FOXA1*, and *FOXA2* in PP2 cells differentiated from hESCs transduced with a scrambled control (SCRAM) or *PDX1* shRNA (sh*PDX1*) lentivirus (*P* = 5.90 x 10^- 5^, 2.74 x 10^-3^, and 0.883 for *PDX1*, *FOXA1*, and *FOXA2*, respectively, in SCRAM compared to sh*PDX1* PP2 cells; student’s t-test, 2-sided; n.s., not significant). (**B**) Percentage of PDX1-bound class I and class II enhancers maintaining or losing FOXA1/2 ChIP-seq peaks in SCRAM or sh*PDX1* PP2. (**C**) Box plots of ATAC-seq and H3K27ac ChIP-seq counts at class I and class II pancreatic enhancers in SCRAM or sh*PDX1* PP2 cells (*P* = < 2.2 x 10^-16^, < 2.2 x 10^-16^, 7.03 x 10^-13^, and < 2.2 x 10^-16^ for SCRAM versus sh*PDX1* of ATAC signal at class I enhancers, ATAC signal at class II enhancers, H3K27ac signal at class I enhancers, and H3K27ac signal at class II enhancers, respectively; Wilcoxon rank sum test, 2-sided). All ChIP-seq and ATAC-seq experiments, n = 2 replicates from independent differentiations.

**Figure S6, related to Figure 6.**
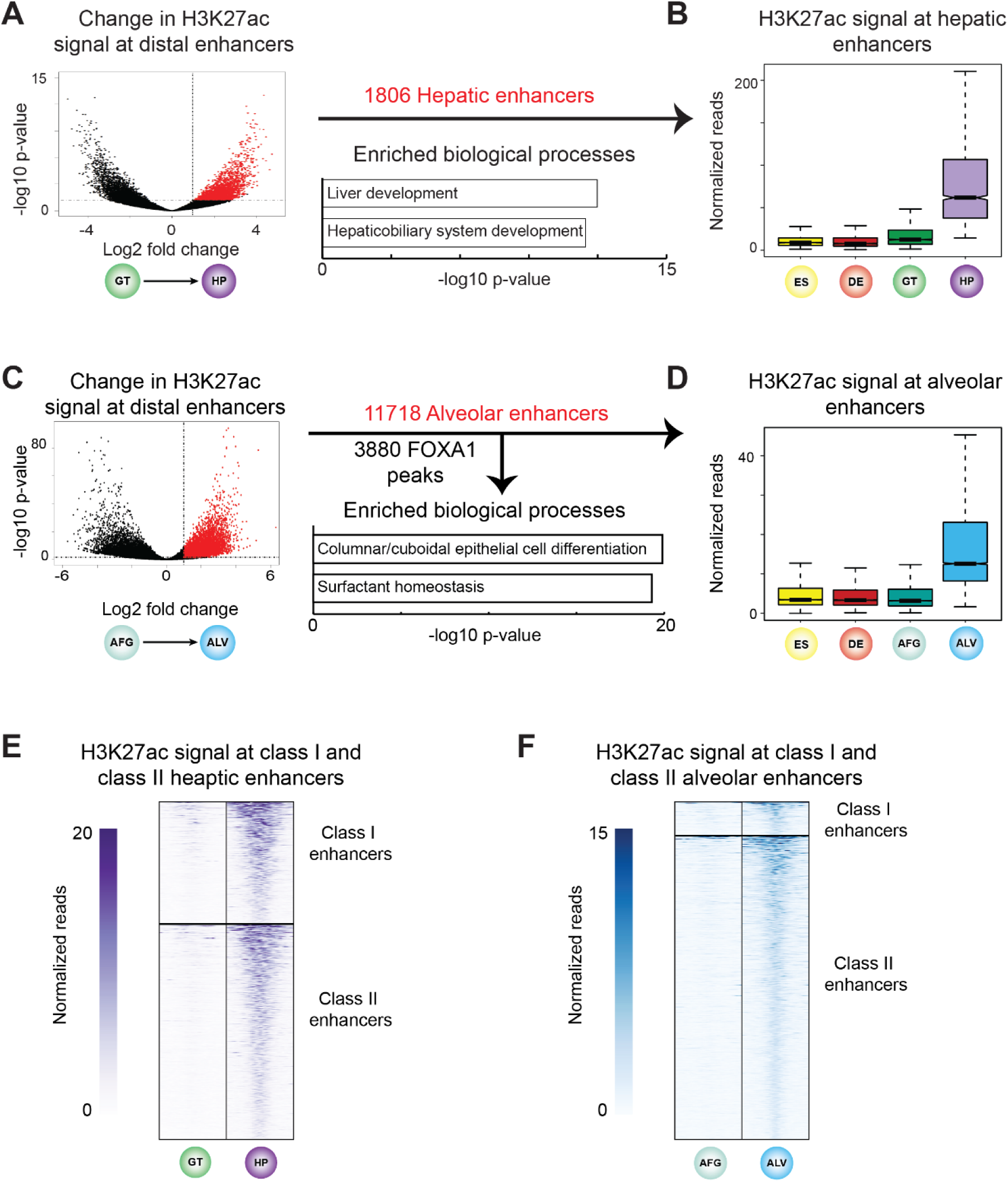
Identification and characterization of hepatic and alveolar enhancers. (**A**) Volcano plot showing identification of hepatic enhancers based on increase in H3K27ac signal from GT to HP (≥ 2-fold increase, *P* adj. < 0.05 at sites > 2.5 kb from TSS). Enriched gene ontology terms of genes linked to hepatic enhancers using GREAT. (**B**) Box plots of H3K27ac ChIP-seq counts at hepatic enhancers. (**C**) Volcano plot showing identification of hepatic enhancers based on increase in H3K27ac signal from AFG to ALV (≥ 2-fold increase, *P* adj. < 0.05 at sites > 2.5 kb from TSS). Enriched gene ontology terms of genes linked to alveolar enhancers using GREAT. (**D**) Box plots of H3K27ac ChIP-seq counts at alveolar enhancers. (**E** and **F**) Heatmaps showing density of H3K27ac ChIP-seq reads at hepatic (**E**) and alveolar (**F**) class I and class II enhancers in GT and HP (**E**) and AFG and ALV (**F**). Heatmaps are centered on H3K27ac peaks and span 5 kb. All ChIP-seq experiments, n = 2 replicates from independent differentiations.

**Figure S7, related to Figure 6.**
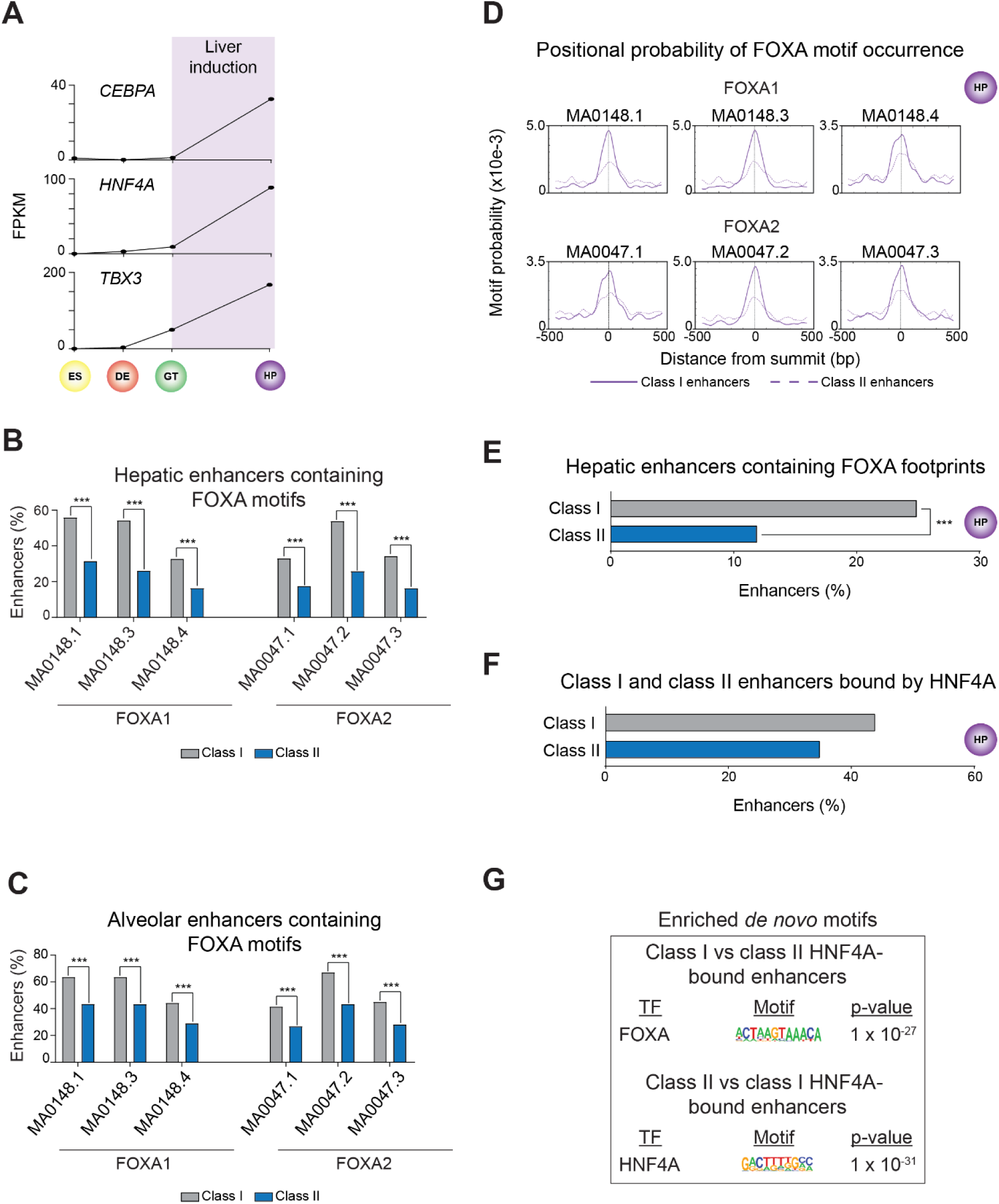
FOXA1/2 binding sites at class I and class II hepatic and alveolar enhancers differ in DNA sequence. (**A**) mRNA expression levels of hepatic transcription factors (TF) determined by RNA-seq. Data are shown as mean fragments per kilobase per million fragments mapped (FPKM) ± S.E.M. in ES, DE, GT (n = 3 independent differentiations), and HP (n = 1 differentiation). (**B** and **C)** Percentage of class I and class II hepatic (**B**) and alveolar (**C**) enhancers with at least one occurrence of selected FOXA1 and FOXA2 motifs (*P* = < 2.2 x 10^-16^, < 2.2 x 10^-16^, 2.42 x 10^-12^, 7.81 x 10^-11^, < 2.2 x 10^-16^, and 1.32 x 10^-14^ for comparisons of occurrences of MA0148.1, MA0148.3, MA0148.4, MA0047.1, MA0047.2, and MA0047.3, respectively, at hepatic enhancers. *P* = 4.88 x 10^-13^, 2.11 x 10^-13^, 6.17 x 10^-9^, 8.89 x 10^-9^, 2.2 x 10^-16^, and 9.75 x 10^-11^ for comparisons of occurrences of MA0148.1, MA0148.3, MA0148.4, MA0047.1, MA0047.2, and MA0047.3, respectively, at alveolar enhancers. Fisher’s exact test, 2-sided). (**D**) Probability (occurrences per base pair) of FOXA1 (MA0148.1, MA0148.3, MA0148.4) and FOXA2 (MA0047.1, MA0047.2, MA0047.3) motifs relative to ATAC-seq peak summits at class I (solid line) and class II (dashed line) hepatic enhancers. ATAC-seq peak summits at class I enhancers are enriched for occurrences compared to summits at class II enhancers (*P* = 4.3 x 10^-14^, 3.3 x 10^-19^, 2.0 x 10^-3^, 5.5 x 10^- 4^, 6.7 x 10^-17^, and 2.1 x 10^-5^ for MA0148.1, MA0148.3, MA0148.4, MA0047.1, MA0047.2, and MA0047.3, respectively; Fisher’s exact test, 1-sided). (**E**) Percentage of hepatic class I and class II enhancers containing FOXA TF ATAC-seq footprints in HP (*P* = 1.01 x 10^-10^ for comparison of class I and class II enhancers; Fisher’s exact test, 2-sided). (**F**) Percentage of hepatic class I and class II enhancers overlapping HNF4A ChIP-seq peaks (within 100 bp from peak) in HP. (**G**) Enriched *de novo* TF binding motifs at HNF4A-bound class I against a background of HNF4A-bound class II enhancers and vice versa. Fisher’s exact test, 1-sided, corrected for multiple comparisons. All ChIP-seq experiments, n = 2 replicates from independent differentiations.

## Supplemental Tables

**Table S1- Related to Figure 1. Genes regulated by FOXA1/2.** (**A**) Genes regulated by FOXA1/2 at GT stage. (**B**) Genes regulated by FOXA1/2 at PP2 stage.

(supplied as Excel file: Table_S1.xlsx)

**Table S2- Related to Figure 2. FOXA TF association with pancreatic enhancers.** (**A**) Class I pancreatic enhancers. (**B**) Class II pancreatic enhancers. (**C**) Class III pancreatic enhancers.

(supplied as Excel file: Table_S2.xlsx)

**Table S3- Related to Figure 3. Characterization of pancreatic enhancers.** (**A**) Pancreatic super-enhancers. (**B**) Chromatin loops at PP2 stage. (**C**) Expressed genes in PP2 proximal to class I pancreatic enhancers. (**D**) Expressed genes in PP2 proximal to class II pancreatic enhancers.

(supplied as Excel file: Table_S3.xlsx)

**Table S4- Related to Figure 4. Differentially enriched de novo motifs between class I and class II pancreatic enhancers.** (**A**) *de novo* motifs enriched in class I pancreatic enhancers over class II pancreatic enhancers. (**B**) *de novo* motifs enriched in class II pancreatic enhancers over class I pancreatic enhancers. (**C**) *de novo* motifs enriched in class I pancreatic enhancers overlapping HNF6 binding sites at PP2 stage compared to class II pancreatic enhancers overlapping HNF6 binding sites at PP2 stage. (**D**) *de novo* motifs enriched in class II pancreatic enhancers overlapping HNF6 binding sites at PP2 stage compared to class I pancreatic enhancers overlapping HNF6 binding sites at PP2 stage. (**E**) *de novo* motifs enriched in class I pancreatic enhancers overlapping PDX1 binding sites at PP2 stage compared to class II pancreatic enhancers overlapping PDX1 binding sites at PP2 stage. (**F**) *de novo* motifs enriched in class II pancreatic enhancers overlapping PDX1 binding sites at PP2 stage compared to class I pancreatic enhancers overlapping PDX1 binding sites at PP2 stage. (**G**) *de novo* motifs enriched in class I pancreatic enhancers overlapping SOX9 binding sites at PP2 stage compared to class II pancreatic enhancers overlapping SOX9 binding sites at PP2 stage. (**H**) *de novo* motifs enriched in class II pancreatic enhancers overlapping SOX9 binding sites at PP2 stage compared to class I pancreatic enhancers overlapping SOX9 binding sites at PP2 stage. (supplied as Excel file: Table_S4.xlsx)

**Table S5- Related to Figure 6. FOXA TF association with hepatic and alveolar enhancers.** (**A**) Class I hepatic enhancers. (**B**) Class II hepatic enhancers. (**C**) Class III hepatic enhancers. (**D**) Class I alveolar enhancers. (**E**) Class II alveolar enhancers. (**F**) Class III alveolar enhancers.

(supplied as Excel file: Table_S5.xlsx)

**Table S6- Related to Figure 6. Differentially enriched de novo motifs between class I and class II hepatic and alveolar enhancers.** (**A**) *de novo* motifs enriched in class I hepatic enhancers over class II hepatic enhancers. (**B**) *de novo* motifs enriched in class II hepatic enhancers over class I hepatic enhancers. (**C**) *de novo* motifs enriched in class I alveolar enhancers over class II alveolar enhancers. (**D**) *de novo* motifs enriched in class II alveolar enhancers over class I alveolar enhancers. (**E**) *de novo* motifs enriched in class I hepatic enhancers overlapping HNF4A binding sites at HP stage compared to class II hepatic enhancers overlapping HNF4A binding sites at HP stage. (**F**) *de novo* motifs enriched in class II hepatic enhancers overlapping HNF4A binding sites at HP stage compared to class I hepatic enhancers overlapping HNF4A binding sites at HP stage. (supplied as Excel file: Table_S6.xlsx)

**Table S7- Related to Figure 7. Differentially enriched known motifs between class I and class II pancreatic, hepatic, and alveolar enhancers.** (**A**) known motifs enriched in class I pancreatic over class I hepatic and alveolar enhancers. (**B**) known motifs enriched in class I hepatic over class I pancreatic and alveolar enhancers. (**C**) known motifs enriched in class I alveolar over class I pancreatic and hepatic enhancers. (**D**) known motifs enriched in class II pancreatic over class II hepatic and alveolar enhancers. (**E**) known motifs enriched in class II hepatic over class II pancreatic and alveolar enhancers. (**F**) known motifs enriched in class II alveolar over class II pancreatic and hepatic enhancers.

(supplied as Excel file: Table_S7.xlsx)

## REFERENCES

Bailey, T.L., and Machanick, P. (2012). Inferring direct DNA binding from ChIP-seq. Nucleic Acids Res 40, e128.

Bonifer, C., and Cockerill, P.N. (2017). Chromatin priming of genes in development: Concepts, mechanisms and consequences. Exp Hematol 49, 1–8.

Buenrostro, J.D., Giresi, P.G., Zaba, L.C., Chang, H.Y., and Greenleaf, W.J. (2013). Transposition of native chromatin for fast and sensitive epigenomic profiling of open chromatin, DNA-binding proteins and nucleosome position. Nat Methods 10, 1213–1218.

Caizzi, L., Ferrero, G., Cutrupi, S., Cordero, F., Ballare, C., Miano, V., Reineri, S., Ricci, L., Friard, O., Testori, A., et al. (2014). Genome-wide activity of unliganded estrogen receptor-alpha in breast cancer cells. Proc Natl Acad Sci U S A 111, 4892–4897.

Carroll, J.S., Liu, X.S., Brodsky, A.S., Li, W., Meyer, C.A., Szary, A.J., Eeckhoute, J., Shao, W., Hestermann, E.V., Geistlinger, T.R., et al. (2005). Chromosome-wide mapping of estrogen receptor binding reveals long-range regulation requiring the forkhead protein FoxA1. Cell 122, 33–43.

Cebola, I., Rodriguez-Segui, S.A., Cho, C.H., Bessa, J., Rovira, M., Luengo, M., Chhatriwala, M., Berry, A., Ponsa-Cobas, J., Maestro, M.A., et al. (2015). TEAD and YAP regulate the enhancer network of human embryonic pancreatic progenitors. Nat Cell Biol 17, 615–626.

Cirillo, L.A., Lin, F.R., Cuesta, I., Friedman, D., Jarnik, M., and Zaret, K.S. (2002). Opening of compacted chromatin by early developmental transcription factors HNF3 (FoxA) and GATA-4. Molecular cell 9, 279–289.

Corso-Diaz, X., de Leeuw, C.N., Alonso, V., Melchers, D., Wong, B.K., Houtman, R., and Simpson, E.M. (2016). Co-activator candidate interactions for orphan nuclear receptor NR2E1. BMC Genomics 17, 832.

Creyghton, M.P., Cheng, A.W., Welstead, G.G., Kooistra, T., Carey, B.W., Steine, E.J., Hanna, J., Lodato, M.A., Frampton, G.M., Sharp, P.A., et al. (2010). Histone H3K27ac separates active from poised enhancers and predicts developmental state. Proc Natl Acad Sci U S A 107, 21931–21936.

Crocker, J., Abe, N., Rinaldi, L., McGregor, A.P., Frankel, N., Wang, S., Alsawadi, A., Valenti, P., Plaza, S., Payre, F., et al. (2015). Low affinity binding site clusters confer hox specificity and regulatory robustness. Cell 160, 191–203.

Dekker, J., Belmont, A.S., Guttman, M., Leshyk, V.O., Lis, J.T., Lomvardas, S., Mirny, L.A., O’Shea, C.C., Park, P.J., Ren, B., et al. (2017). The 4D nucleome project. Nature 549, 219–226.

Deng, X., Zhang, X., Li, W., Feng, R.X., Li, L., Yi, G.R., Zhang, X.N., Yin, C., Yu, H.Y., Zhang, J.P., et al. (2018). Chronic Liver Injury Induces Conversion of Biliary Epithelial Cells into Hepatocytes. Cell Stem Cell 23, 114–122 e113.

Dixon, J.R., Jung, I., Selvaraj, S., Shen, Y., Antosiewicz-Bourget, J.E., Lee, A.Y., Ye, Z., Kim, A., Rajagopal, N., Xie, W., et al. (2015). Chromatin architecture reorganization during stem cell differentiation. Nature 518, 331–336.

Dobin, A., Davis, C.A., Schlesinger, F., Drenkow, J., Zaleski, C., Jha, S., Batut, P., Chaisson, M., and Gingeras, T.R. (2013). STAR: ultrafast universal RNA-seq aligner. Bioinformatics 29, 15–21.

Donaghey, J., Thakurela, S., Charlton, J., Chen, J.S., Smith, Z.D., Gu, H., Pop, R., Clement, K., Stamenova, E.K., Karnik, R., et al. (2018). Genetic determinants and epigenetic effects of pioneer-factor occupancy. Nat Genet 50, 250–258.

Durand, N.C., Robinson, J.T., Shamim, M.S., Machol, I., Mesirov, J.P., Lander, E.S., and Aiden, E.L. (2016). Juicebox Provides a Visualization System for Hi-C Contact Maps with Unlimited Zoom. Cell Syst 3, 99–101.

Farley, E.K., Olson, K.M., Zhang, W., Brandt, A.J., Rokhsar, D.S., and Levine, M.S. (2015). Suboptimization of developmental enhancers. Science 350, 325–328.

Fornes, O., Castro-Mondragon, J.A., Khan, A., van der Lee, R., Zhang, X., Richmond, P.A., Modi, B.P., Correard, S., Gheorghe, M., Baranasic, D., et al. (2020). JASPAR 2020: update of the open-access database of transcription factor binding profiles. Nucleic Acids Res 48, D87–D92.

Genga, R.M.J., Kernfeld, E.M., Parsi, K.M., Parsons, T.J., Ziller, M.J., and Maehr, R. (2019). Single-Cell RNA-Sequencing-Based CRISPRi Screening Resolves Molecular Drivers of Early Human Endoderm Development. Cell Rep 27, 708–718 e710.

Grant, C.E., Bailey, T.L., and Noble, W.S. (2011). FIMO: scanning for occurrences of a given motif. Bioinformatics 27, 1017–1018.

Gualdi, R., Bossard, P., Zheng, M., Hamada, Y., Coleman, J.R., and Zaret, K.S. (1996). Hepatic specification of the gut endoderm in vitro: cell signaling and transcriptional control. Genes Dev 10, 1670–1682.

Hawkins, F., Kramer, P., Jacob, A., Driver, I., Thomas, D.C., McCauley, K.B., Skvir, N., Crane, A.M., Kurmann, A.A., Hollenberg, A.N., et al. (2017). Prospective isolation of NKX2-1-expressing human lung progenitors derived from pluripotent stem cells. J Clin Invest 127, 2277–2294.

Heinz, S., Benner, C., Spann, N., Bertolino, E., Lin, Y.C., Laslo, P., Cheng, J.X., Murre, C., Singh, H., and Glass, C.K. (2010). Simple combinations of lineage-determining transcription factors prime cis-regulatory elements required for macrophage and B cell identities. Molecular cell 38, 576–589.

Herriges, M., and Morrisey, E.E. (2014). Lung development: orchestrating the generation and regeneration of a complex organ. Development 141, 502–513.

Huang, W., Wang, G., Delaspre, F., Vitery Mdel, C., Beer, R.L., and Parsons, M.J. (2014). Retinoic acid plays an evolutionarily conserved and biphasic role in pancreas development. Dev Biol 394, 83–93.

Hurley, K., Ding, J., Villacorta-Martin, C., Herriges, M.J., Jacob, A., Vedaie, M., Alysandratos, K.D., Sun, Y.L., Lin, C., Werder, R.B., et al. (2020). Reconstructed Single-Cell Fate Trajectories Define Lineage Plasticity Windows during Differentiation of Human PSC-Derived Distal Lung Progenitors. Cell Stem Cell 26, 593–608 e598.

Hurtado, A., Holmes, K.A., Ross-Innes, C.S., Schmidt, D., and Carroll, J.S. (2011). FOXA1 is a key determinant of estrogen receptor function and endocrine response. Nat Genet 43, 27–33.

Ikonomou, L., Herriges, M.J., Lewandowski, S.L., Marsland, R., 3rd, Villacorta-Martin, C., Caballero, I.S., Frank, D.B., Sanghrajka, R.M., Dame, K., Kandula, M.M., et al. (2020). The in vivo genetic program of murine primordial lung epithelial progenitors. Nat Commun 11, 635.

Iwafuchi, M., Cuesta, I., Donahue, G., Takenaka, N., Osipovich, A.B., Magnuson, M.A., Roder, H., Seeholzer, S.H., Santisteban, P., and Zaret, K.S. (2020). Gene network transitions in embryos depend upon interactions between a pioneer transcription factor and core histones. Nat Genet 52, 418–427.

Iwafuchi-Doi, M., Donahue, G., Kakumanu, A., Watts, J.A., Mahony, S., Pugh, B.F., Lee, D., Kaestner, K.H., and Zaret, K.S. (2016). The Pioneer Transcription Factor FoxA Maintains an Accessible Nucleosome Configuration at Enhancers for Tissue-Specific Gene Activation. Molecular cell 62, 79–91.

Jacob, A., Morley, M., Hawkins, F., McCauley, K.B., Jean, J.C., Heins, H., Na, C.L., Weaver, T.E., Vedaie, M., Hurley, K., et al. (2017). Differentiation of Human Pluripotent Stem Cells into Functional Lung Alveolar Epithelial Cells. Cell Stem Cell 21, 472–488 e410.

Jacob, A., Vedaie, M., Roberts, D.A., Thomas, D.C., Villacorta-Martin, C., Alysandratos, K.D., Hawkins, F., and Kotton, D.N. (2019). Derivation of self-renewing lung alveolar epithelial type II cells from human pluripotent stem cells. Nat Protoc 14, 3303–3332.

Jin, W., Mulas, F., Gaertner, B., Sui, Y., Wang, J., Matta, I., Zeng, C., Vinckier, N., Wang, A., Nguyen-Ngoc, K.V., et al. (2019). A Network of microRNAs Acts to Promote Cell Cycle Exit and Differentiation of Human Pancreatic Endocrine Cells. iScience 21, 681–694.

Kent, W.J., Sugnet, C.W., Furey, T.S., Roskin, K.M., Pringle, T.H., Zahler, A.M., and Haussler, D. (2002). The human genome browser at UCSC. Genome Res 12, 996–1006.

Kheolamai, P., and Dickson, A.J. (2009). Liver-enriched transcription factors are critical for the expression of hepatocyte marker genes in mES-derived hepatocyte-lineage cells. BMC Mol Biol 10, 35.

Kobberup, S., Nyeng, P., Juhl, K., Hutton, J., and Jensen, J. (2007). ETS-family genes in pancreatic development. Dev Dyn 236, 3100–3110.

Lee, C.S., Friedman, J.R., Fulmer, J.T., and Kaestner, K.H. (2005). The initiation of liver development is dependent on Foxa transcription factors. Nature 435, 944–947.

Lee, K., Cho, H., Rickert, R.W., Li, Q.V., Pulecio, J., Leslie, C.S., and Huangfu, D. (2019). FOXA2 Is Required for Enhancer Priming during Pancreatic Differentiation. Cell Rep 28, 382–393 e387.

Li, H., and Durbin, R. (2009). Fast and accurate short read alignment with Burrows-Wheeler transform. Bioinformatics 25, 1754–1760.

Li, H., Handsaker, B., Wysoker, A., Fennell, T., Ruan, J., Homer, N., Marth, G., Abecasis, G., Durbin, R., and Genome Project Data Processing, S. (2009). The Sequence Alignment/Map format and SAMtools. Bioinformatics 25, 2078–2079.

Love, M.I., Huber, W., and Anders, S. (2014). Moderated estimation of fold change and dispersion for RNA-seq data with DESeq2. Genome Biol 15, 550.

Loven, J., Hoke, H.A., Lin, C.Y., Lau, A., Orlando, D.A., Vakoc, C.R., Bradner, J.E., Lee, T.I., and Young, R.A. (2013). Selective inhibition of tumor oncogenes by disruption of super-enhancers. Cell 153, 320–334.

Lupien, M., Eeckhoute, J., Meyer, C.A., Wang, Q., Zhang, Y., Li, W., Carroll, J.S., Liu, X.S., and Brown, M. (2008). FoxA1 translates epigenetic signatures into enhancer-driven lineage-specific transcription. Cell 132, 958–970.

Maddalo, D., Manchado, E., Concepcion, C.P., Bonetti, C., Vidigal, J.A., Han, Y.C., Ogrodowski, P., Crippa, A., Rekhtman, N., de Stanchina, E., et al. (2014). In vivo engineering of oncogenic chromosomal rearrangements with the CRISPR/Cas9 system. Nature 516, 423–427.

Magnani, L., and Lupien, M. (2014). Chromatin and epigenetic determinants of estrogen receptor alpha (ESR1) signaling. Mol Cell Endocrinol 382, 633–641.

Mamidi, A., Prawiro, C., Seymour, P.A., de Lichtenberg, K.H., Jackson, A., Serup, P., and Semb, H. (2018). Mechanosignalling via integrins directs fate decisions of pancreatic progenitors. Nature 564, 114–118.

McLean, C.Y., Bristor, D., Hiller, M., Clarke, S.L., Schaar, B.T., Lowe, C.B., Wenger, A.M., and Bejerano, G. (2010). GREAT improves functional interpretation of cis-regulatory regions. Nature biotechnology 28, 495–501.

Moffat, J., Grueneberg, D.A., Yang, X., Kim, S.Y., Kloepfer, A.M., Hinkle, G., Piqani, B., Eisenhaure, T.M., Luo, B., Grenier, J.K., et al. (2006). A lentiviral RNAi library for human and mouse genes applied to an arrayed viral high-content screen. Cell 124, 1283–1298.

Paakinaho, V., Swinstead, E.E., Presman, D.M., Grontved, L., and Hager, G.L. (2019). Meta-analysis of Chromatin Programming by Steroid Receptors. Cell Rep 28, 3523–3534 e3522.

Papaioannou, V.E. (2014). The T-box gene family: emerging roles in development, stem cells and cancer. Development 141, 3819–3833.

Quinlan, A.R., and Hall, I.M. (2010). BEDTools: a flexible suite of utilities for comparing genomic features. Bioinformatics 26, 841–842.

Rada-Iglesias, A., Bajpai, R., Swigut, T., Brugmann, S.A., Flynn, R.A., and Wysocka, J. (2011). A unique chromatin signature uncovers early developmental enhancers in humans. Nature 470, 279–283.

Ramana, C.V. (2019). Insights into the Signal Transduction Pathways of Mouse Lung Type II Cells Revealed by Transcription Factor Profiling in the Transcriptome. Genomics Inform 17, e8.

Ramirez, F., Ryan, D.P., Gruning, B., Bhardwaj, V., Kilpert, F., Richter, A.S., Heyne, S., Dundar, F., and Manke, T. (2016). deepTools2: a next generation web server for deep-sequencing data analysis. Nucleic Acids Res 44, W160–165.

Rankin, S.A., McCracken, K.W., Luedeke, D.M., Han, L., Wells, J.M., Shannon, J.M., and Zorn, A.M. (2018). Timing is everything: Reiterative Wnt, BMP and RA signaling regulate developmental competence during endoderm organogenesis. Dev Biol 434, 121–132.

Rao, S.S., Huntley, M.H., Durand, N.C., Stamenova, E.K., Bochkov, I.D., Robinson, J.T., Sanborn, A.L., Machol, I., Omer, A.D., Lander, E.S., et al. (2014). A 3D map of the human genome at kilobase resolution reveals principles of chromatin looping. Cell 159, 1665–1680.

Rezania, A., Bruin, J.E., Arora, P., Rubin, A., Batushansky, I., Asadi, A., O’Dwyer, S., Quiskamp, N., Mojibian, M., Albrecht, T., et al. (2014). Reversal of diabetes with insulin-producing cells derived in vitro from human pluripotent stem cells. Nature biotechnology 32, 1121–1133.

Schulz, T.C., Young, H.Y., Agulnick, A.D., Babin, M.J., Baetge, E.E., Bang, A.G., Bhoumik, A., Cepa, I., Cesario, R.M., Haakmeester, C., et al. (2012). A scalable system for production of functional pancreatic progenitors from human embryonic stem cells. PLoS One 7, e37004.

Sekiya, T., Muthurajan, U.M., Luger, K., Tulin, A.V., and Zaret, K.S. (2009). Nucleosome-binding affinity as a primary determinant of the nuclear mobility of the pioneer transcription factor FoxA. Genes Dev 23, 804–809.

Soufi, A., Garcia, M.F., Jaroszewicz, A., Osman, N., Pellegrini, M., and Zaret, K.S. (2015). Pioneer transcription factors target partial DNA motifs on nucleosomes to initiate reprogramming. Cell 161, 555–568.

Sung, M.H., Guertin, M.J., Baek, S., and Hager, G.L. (2014). DNase footprint signatures are dictated by factor dynamics and DNA sequence. Molecular cell 56, 275–285.

Swinstead, E.E., Miranda, T.B., Paakinaho, V., Baek, S., Goldstein, I., Hawkins, M., Karpova, T.S., Ball, D., Mazza, D., Lavis, L.D., et al. (2016). Steroid Receptors Reprogram FoxA1 Occupancy through Dynamic Chromatin Transitions. Cell 165, 593–605.

Tata, P.R., Chow, R.D., Saladi, S.V., Tata, A., Konkimalla, A., Bara, A., Montoro, D., Hariri, L.P., Shih, A.R., Mino-Kenudson, M., et al. (2018). Developmental History Provides a Roadmap for the Emergence of Tumor Plasticity. Dev Cell 44, 679–693 e675.

Trapnell, C., Williams, B.A., Pertea, G., Mortazavi, A., Kwan, G., van Baren, M.J., Salzberg, S.L., Wold, B.J., and Pachter, L. (2010). Transcript assembly and quantification by RNA-Seq reveals unannotated transcripts and isoform switching during cell differentiation. Nature biotechnology 28, 511–515.

Tuteja, G., Jensen, S.T., White, P., and Kaestner, K.H. (2008). Cis-regulatory modules in the mammalian liver: composition depends on strength of Foxa2 consensus site. Nucleic Acids Res 36, 4149–4157.

Wan, H., Dingle, S., Xu, Y., Besnard, V., Kaestner, K.H., Ang, S.L., Wert, S., Stahlman, M.T., and Whitsett, J.A. (2005). Compensatory roles of Foxa1 and Foxa2 during lung morphogenesis. J Biol Chem 280, 13809–13816.

Wang, A., Yue, F., Li, Y., Xie, R., Harper, T., Patel, N.A., Muth, K., Palmer, J., Qiu, Y., Wang, J., et al. (2015). Epigenetic priming of enhancers predicts developmental competence of hESC-derived endodermal lineage intermediates. Cell Stem Cell 16, 386–399.

Wang, Q., Li, W., Zhang, Y., Yuan, X., Xu, K., Yu, J., Chen, Z., Beroukhim, R., Wang, H., Lupien, M., et al. (2009). Androgen receptor regulates a distinct transcription program in androgen-independent prostate cancer. Cell 138, 245–256.

Whyte, W.A., Orlando, D.A., Hnisz, D., Abraham, B.J., Lin, C.Y., Kagey, M.H., Rahl, P.B., Lee, T.I., and Young, R.A. (2013). Master transcription factors and mediator establish super-enhancers at key cell identity genes. Cell 153, 307–319.

Xie, R., Everett, L.J., Lim, H.W., Patel, N.A., Schug, J., Kroon, E., Kelly, O.G., Wang, A., D’Amour, K.A., Robins, A.J., et al. (2013). Dynamic chromatin remodeling mediated by polycomb proteins orchestrates pancreatic differentiation of human embryonic stem cells. Cell Stem Cell 12, 224–237.

Zhang, Y., Fang, B., Emmett, M.J., Damle, M., Sun, Z., Feng, D., Armour, S.M., Remsberg, J.R., Jager, J., Soccio, R.E., et al. (2015). GENE REGULATION. Discrete functions of nuclear receptor Rev-erbalpha couple metabolism to the clock. Science 348, 1488–1492.

Zhang, Y., Liu, T., Meyer, C.A., Eeckhoute, J., Johnson, D.S., Bernstein, B.E., Nusbaum, C., Myers, R.M., Brown, M., Li, W., et al. (2008). Model-based analysis of ChIP-Seq (MACS). Genome Biol 9, R137.

Zhou, Y., Zhou, B., Pache, L., Chang, M., Khodabakhshi, A.H., Tanaseichuk, O., Benner, C., and Chanda, S.K. (2019). Metascape provides a biologist-oriented resource for the analysis of systems-level datasets. Nat Commun 10, 1523.

Zorn, A.M., and Wells, J.M. (2009). Vertebrate endoderm development and organ formation. Annu Rev Cell Dev Biol 25, 221–251.

